# Predicting Functional Conformational Ensembles and Binding Mechanisms of Convergent Evolution for SARS-CoV-2 Spike Omicron Variants Using AlphaFold2 Sequence Scanning Adaptations and Molecular Dynamics Simulations

**DOI:** 10.1101/2024.04.02.587850

**Authors:** Nishank Raisinghani, Mohammed Alshahrani, Grace Gupta, Sian Xiao, Peng Tao, Gennady Verkhivker

## Abstract

In this study, we combined AlphaFold-based approaches for atomistic modeling of multiple protein states and microsecond molecular simulations to accurately characterize conformational ensembles and binding mechanisms of convergent evolution for the SARS-CoV-2 Spike Omicron variants BA.1, BA.2, BA.2.75, BA.3, BA.4/BA.5 and BQ.1.1. We employed and validated several different adaptations of the AlphaFold methodology for modeling of conformational ensembles including the introduced randomized full sequence scanning for manipulation of sequence variations to systematically explore conformational dynamics of Omicron Spike protein complexes with the ACE2 receptor. Microsecond atomistic molecular dynamic simulations provide a detailed characterization of the conformational landscapes and thermodynamic stability of the Omicron variant complexes. By integrating the predictions of conformational ensembles from different AlphaFold adaptations and applying statistical confidence metrics we can expand characterization of the conformational ensembles and identify functional protein conformations that determine the equilibrium dynamics for the Omicron Spike complexes with the ACE2. Conformational ensembles of the Omicron RBD-ACE2 complexes obtained using AlphaFold-based approaches for modeling protein states and molecular dynamics simulations are employed for accurate comparative prediction of the binding energetics revealing an excellent agreement with the experimental data. In particular, the results demonstrated that AlphaFold-generated extended conformational ensembles can produce accurate binding energies for the Omicron RBD-ACE2 complexes. The results of this study suggested complementarities and potential synergies between AlphaFold predictions of protein conformational ensembles and molecular dynamics simulations showing that integrating information from both methods can potentially yield a more adequate characterization of the conformational landscapes for the Omicron RBD-ACE2 complexes. This study provides insights in the interplay between conformational dynamics and binding, showing that evolution of Omicron variants through acquisition of convergent mutational sites may leverage conformational adaptability and dynamic couplings between key binding energy hotspots to optimize ACE2 binding affinity and enable immune evasion.

## Introduction

The comprehensive array of structural and biochemical investigations focused on the Spike (S) glycoprotein of the SARS-CoV-2 virus has provided crucial understanding of the mechanisms dictating virus transmission and evasion of the immune system. This glycoprotein serves as the gateway for viral entry into host cells and undergoes significant conformational alterations, transitioning between closed and open states. These transitions are orchestrated by the flexible amino (N)-terminal S1 subunit, which encompasses the N-terminal domain (NTD), the receptor-binding domain (RBD), and two structurally conserved subdomains—SD1 and SD2. ^1–9^ The dynamic interplay among these structural domains is crucial for regulating conformational transitions within the S protein, facilitating shifts between the RBD-down closed state and the RBD-up open state, consequently enabling a variety of functional responses. Additionally, the functional motions of the NTD, RBD, SD1, and SD2 subdomains are synchronized as the S1 subunit undergoes global movements. These coordinated movements are facilitated through long-range communications with the structurally rigid S2 subunit.^10–15^ The synergistic interplay of functional motions within both S1 and S2 is paramount in mediating crucial interactions between the S protein and the host cell receptor ACE2. Moreover, it governs a wide array of interactions between the S protein and various classes of antibodies, thereby impacting the host immune responses triggered by the virus. The ability of the S protein to stochastically sample distinct structural states is fundamental for its effectiveness and specificity in recognizing host cell receptors and evading immune detection. The abundance of data and insights gained from biophysical studies has enriched our understanding of the S protein trimer, illuminating the complex interplay between thermodynamics and kinetics that govern spike mechanisms. These studies have revealed how mutations and long-range interactions between the dynamic S1 subunit and the more rigid S2 subunit orchestrate coordinated structural alterations within the S protein trimer and control population shifts between the open and closed RBD states, thus modulating interactions with various binding partners and shaping immune responses.^16–18^

The growing accessibility of cryo-electron microscopy (cryo-EM) and X-ray structures for the S protein variants of concern (VOCs) has significantly broadened our understanding of the evolutionary adaptability of the S protein and the diverse array of molecular mechanisms involved. These structural studies have revealed a multitude of functional states of the S protein and its interactions with antibodies, contributing to a complex and adaptable dynamic landscape with a diverse range of binding epitopes.^19–28^ The cryo-EM structures and biochemical analyses of the S trimers of different subvariants emerged during Omicron evolution including BA.1, BA.2, BA.3, and BA.4/BA.5, have revealed significant similarities and subtle differences in binding energetics with the host receptor. The binding affinities of Omicron BA.2 with ACE2 were observed to be stronger than those of BA.3 and BA.1, as indicated by the structures of the RBD-ACE2 complexes for the BA.1.1, BA.2, and BA.3 variants.^29^ Additionally, the Omicron BA.2 trimer exhibited a higher ACE2 binding affinity compared to both the S Wu-Hu-1 trimer and the S Omicron BA.1 trimer.^30^ The surface plasmon resonance (SPR) results showed that the Omicron BA.4/5 RBD had only a slightly higher binding affinity for ACE2 than the ancestral Wu-Hu-1 strain and the BA.1 variants.^31^ The biochemical analyses of BA.1, BA.2, BA.3, and BA.4/BA.5 variants demonstrated higher binding affinities for BA.2 compared to other Omicron variants.^32,33^ Biophysical studies of the Omicron BA.2.75 subvariant demonstrated a remarkable 9-fold enhancement in the binding affinity with ACE2 compared to its parental BA.2 variant, establishing it as having the strongest ACE2 binding among all S variants.^34^ Additionally, cryo-EM conformations of the BA.2.75 S trimer and structures of the open BA.2.75 S trimer complexes with ACE2 revealed that the BA.2.75 S-trimer exhibited the highest stability, followed by BA.1, BA.5, and BA.2 variants.^35^ The SPR experiments confirmed these findings, showing that the BA.2.75 subvariant exhibited a notably higher ACE2 binding affinity which is approximately 4-6 times greater than the other Omicron variants.^34–36^ Structural-functional studies of the Omicron BA.1, BA.2, BA.2.12.1, BA.4, and BA.5 subvariants provided further support to the mechanism in which the combined effect of the enhanced ACE2 receptor binding and stronger immune evasion may have contributed to the rapid spread of Omicron sublineages.^37^ Structural and virological analysis of the BA.4/BA.5 variant showed that dissociation constants and the binding affinity of the S RBDs of BA.2 and BA.4/5 are similar where F486V in the BA.4/5 S RBD may decrease the hydrophobic interaction with ACE2, while Q493 of the BA.4/5 S RBD by forming a hydrogen bond with the ACE2 residue H34 can compensate for the loss of binding.^38,39^

BQ.1.1 S protein harbors five convergent substitutions R346T, K444T, L452R, N460K, and F486V. Yeast surface display assays showed that the dissociation constant (*K*_D_) value of BQ.1.1 RBD (0.66□±□0.11) to the human ACE2 molecule is lower than that of the parental BA.5 RBD (1.08□±□0.16) suggesting that BQ.1.1 increased the binding affinity to human ACE2 than BA.5 variant.^39^ Biophysical studies also revealed that BQ.1.1 exhibited stronger antibody evasion owing to convergent mutations R346T and N460K.^40^ Another study indicated that N460K-bearing subvariants BQ.1 and BQ.1.1 exhibit the strongest neutralization resistance compared with BA.4/5 variants suggesting that N460K mutation, and to a lesser extent, R346T, K444T are critical for the enhanced resistance of the BQ.1 and BQ.1.1 subvariants. Structural modeling of BQ.1.1 variant showed the important role of convergent mutational site F486V that can impact binding affinity with both ACE2 and antibodies, whereas the R346T and K444T mutations are likely responsible for evasion of specific class 3 antibody recognition.^41^ X-ray crystallographic analysis of the BQ.1.1 RBD-human ACE2 complex showed that R346T and K444T are not directly involved in the interaction with ACE2 but compared to the BA.5-human ACE2 complex structure may induce a different pattern of the RBD dynamics in the flexible regions.^42^ Functional studies confirmed that emergence of BQ.1.1 bearing R346T along with K444T and N460K substitutions are primarily associated with escape from monoclonal antibodies and vaccine-induced antibodies.^43,44^

The patterns of convergent evolution was analyze based the mutations which emerged at least three times independently in different linages,^45^ suggesting that spatial clustering of amino acids under convergent evolution creates suggest potential for synergistic epistatic effects.^46,47^ Convergent evolution of mutations was also seen in the immune-evasive XBB lineages that harbor F496P and acquired additional mutations including R403K, V445S, L455F, F456L, and K478R allowing for escape from neutralizing antibodies generated through both repeated vaccination and natural infection.^48–50^ The latest structures and SPR-measured binding of S-RBD with human ACE2 for BA.4/BA.5, BQ.1, BQ.1.1, XBB and XBB.1.5 variants showed that BQ.1.1 affinity is comparable to that of BA.4/5 highlighting the key role of F486V as binding hotspot and also suggesting that R346T substitution can enhance ACE2 binding using long-range couplings with Q493 position.^51^ These studies echoed recent deep mutational scanning (DMS) experiments of BQ.1.1 and XBB.1.5 RBDs binding with ACE2 showing epistatic couplings between R493Q and mutations at positions Y453, L455, and F456.^52^ Convergent evolution patterns of S mutations showed that coordination of evolutionary paths at different sites may be due to epistatic rather than random selection of mutations.^53,54^

Computer simulation studies provided important atomistic insights into understanding the dynamics of the SARS-CoV-2 S protein and binding with diverse binding partners.^55–60^ Conformational dynamics and allosteric modulation of SARS-CoV-2 S in the absence or presence of ligands was studied using smFRET imaging assay, showing presence of long-range allosteric control of the RBD equilibrium, which in turn regulates the exposure of the binding site and antibody binding.^61^ Integrative computational modeling approaches revealed that the S protein could function as an allosteric regulatory machinery controlled by stable allosteric hotspots acting as drives drivers and regulators of spike activity.^62–67^ By combining atomistic simulations and a community-based network model of epistatic couplings we found that convergent Omicron mutations can display epistatic relationships with the major stability and binding affinity hotspots which may allow for the observed broad antibody resistance.^65^ Analysis of conformational dynamics and allosteric communications in the Omicron BA.1, BA.2, BA.3 and BA.4/BA.5 complexes with the ACE2 host receptor characterized regions of epistatic couplings that are centered at the binding affinity hotspots N501Y and Q498R enabling accumulation of multiple Omicron immune escape mutations at other sites.^66^ MD simulations and Markov state models systematically characterized conformational landscapes and identify specific dynamic signatures of Omicron variants and their complexes showing that convergent mutation sites could control evolution allosteric pockets through modulation of conformational plasticity in the flexible adaptable regions.^67^ Recent computational studies suggested that Omicron mutations have variant-specific effect on conformational dynamics changes in the S protein including allosterically induced plasticity at the remote regions leading to the formation and evolution of druggable cryptic pockets.^68,69^

Accurate characterization of the structural ensembles is paramount for robust predictions of the S protein activity and binding with the ACE2 and antibodies. The emergence of AlphaFold2 (AF2) technology has revolutionized protein structure modeling in structural biology, representing a significant leap forward.^70,71^ While AF2-based methods have made impressive strides in predicting static protein structures, they encounter hurdles in precisely capturing conformational dynamics, functional protein ensembles, conformational changes, and allosteric states.^72^ Recent studies underscore that while AF2 methods demonstrate proficiency in predicting individual protein structures, expanding this capability to accurately forecast conformational ensembles and map allosteric landscapes still poses a substantial challenge.^73–77^ The limitations of AF2 methods in predicting multiple protein conformations may be linked to the intrinsic training bias towards experimentally determined, thermodynamically stable structures, and MSAs containing evolutionary information used to infer the ground protein states. Efforts to optimize the AF2 methodology for predicting alternative conformational states were primarily focused on manipulating the MSA information to encode coevolutionary signals, aiming to capture not only the most thermodynamically probable protein state but also other functional conformational states of a protein.^73,74^ One approach involves subsampling of MSAs to reduce depth, resulting in shallow MSAs.^73^ This strategy aims to enhance diversity and generate a broader range of AF2 output models, thereby improving the ability to capture experimentally validated alternative conformational states of proteins. The alternative AF2-based approach, SPEACH_AF (Sampling Protein Ensembles and Conformational Heterogeneity with AlphaFold2) altera the MSA through in silico mutagenesis, where specific residues within the MSA are replaced to induce changes.^74^ Recent studies have investigated the combination of shallow MSA with state-annotated templates, which integrate functional or structural properties of GPCRs and protein kinases.^78^ Another AF2 adaption termed AF-Cluster employs a simple method of subsampling MSAs followed by clustering related or functionally similar sequences. This approach allows for the prediction of alternative protein states and has shown promise in identifying previously unknown fold-switched states, which were later validated by NMR analysis.^79^

In this study, we employed an AI-enabled integrative simulation approach for probing conformational ensembles and binding energetics of the Omicron RBD-ACE2 complexes. Several different adaptations of the AF2 methodology along with MD simulations were used for a comparative characterization of structures, conformational ensembles and subsequent computations of binding affinities of the Omicron RBD-ACE2 complexes including BA.1, BA.2, BA.2.75, BA.3, BA.4/BA.5 and BQ.1.1 variants (Figure 1). Structural organization of the RBD-ACE2 complexes for all Omicron variants is remarkably similar and the composition of the binding epitopes across Omicron variants is preserved (Figure 1). Using a combination of the AF2-based adaptations we can efficiently expand characterization of the conformational ensembles for the RBD-ACE2 complexes capturing conformational details of the RBD fold and variant-specific functional adjustments of the RBM loop for binding. We demonstrate that AF2-based statistical assessment of generated models can be used to accurately predict the mutation-induced dynamic changes and determine functional protein ensembles.

**Figure 1.**
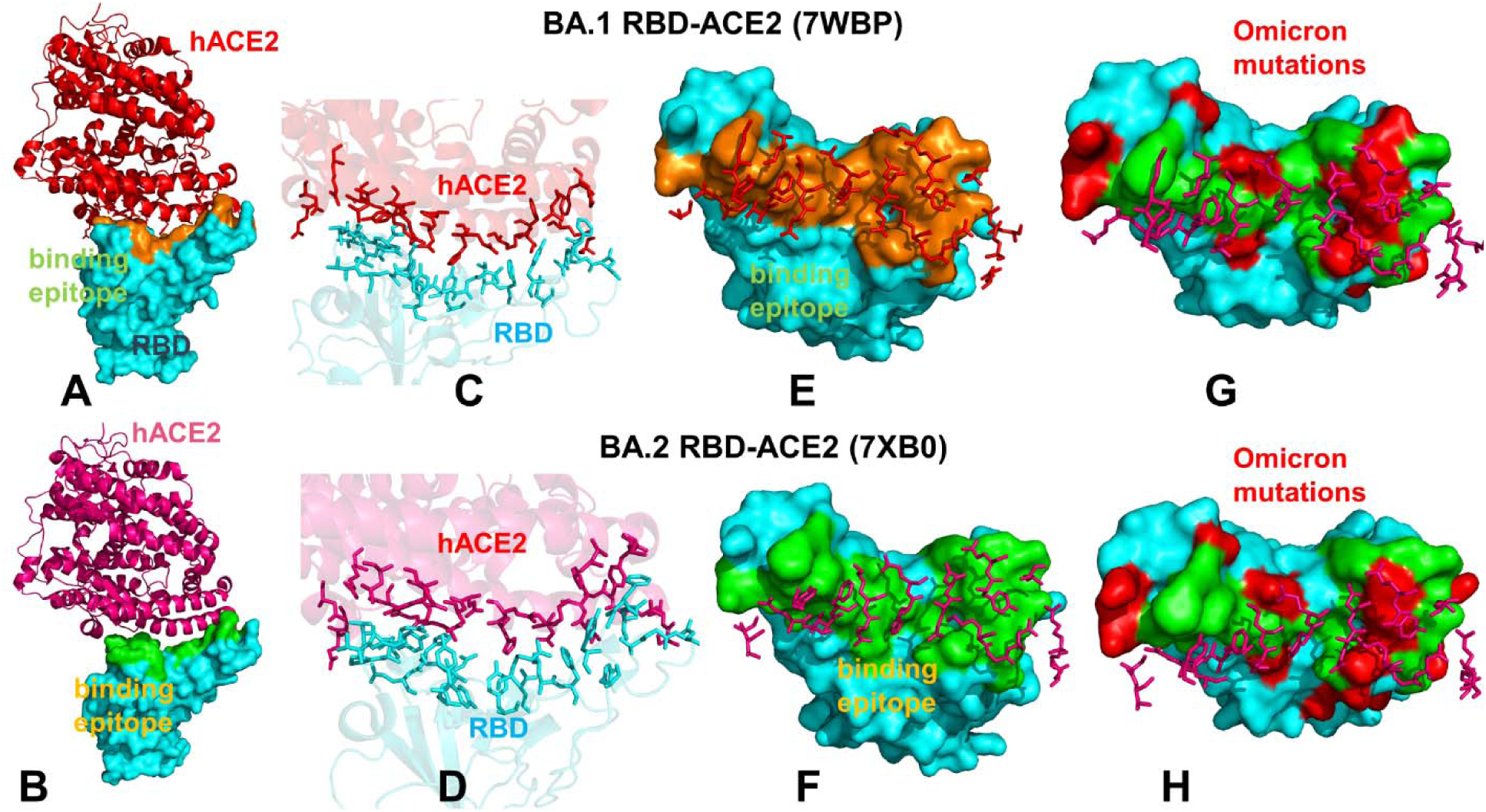
Structural organization and binding epitopes of the SARS-CoV-2-RBD Omicron complexes with human ACE enzyme. The cryo-EM structure of the Omicron RBD BA.1-ACE2 complex (pdb id 7WBP) (A). The RBD is in cyan surface and ACE2 is in red ribbons. (B) The cryo-EM structure of the Omicron RBD BA.2-ACE2 complex (pdb id 7XB0). The RBD is in cyan surface and the ACE2 is in pink ribbons. (C) The binding interface for the BA.1 RBD-ACE2 complex. The RBD binding residues are in cyan sticks, ACE2 binding residues are in red sticks. (D) The biding interface for the BA.2 RBD-ACE2 complex. The RBD binding residues are in cyan sticks, ACE2 binding residues are in pink sticks. (E) The binding epitope for the BA.1 RBD-ACE2 complex. The RBD-BA.1 binding epitope is in orange surface. The ACE2 binding residues are shown in pink sticks. (F) The binding epitope for the BA.2 RBD-ACE2 complex. The RBD-BA.1 binding epitope is in green surface. The ACE2 binding residues are shown in pink sticks. (G) The binding epitope residues (in green surface) and BA.1 RBD Omicron mutations (in red surface). The RBD is shown in cyan surface. (H) The map of the binding epitope residues (in green surface) and BA.2 RBD Omicron mutations (in red surface). Microsecond atomistic simulations are also conducted to provide a detailed characterization of the conformational landscapes and thermodynamic stability of the Omicron variant complexes. We find that AF2 predictions of structural ensembles are consistent with the conformational flexibility patterns revealed in atomistic MD simulations. We leverage AF2-based structural ensembles and MD-generated equilibrium ensembles for a comparative prediction of the binding energetics and affinities for the Omicron RBD-ACE2 complexes, resulting in remarkably similar trends and strong correlation with the experimental data. This integrative analysis shows that combining MD simulations together with AF2-based prediction of conformational ensembles could provide more comprehensive view of the conformational landscapes and binding mechanisms for Omicron variants. We also examine how evolution of Omicron variants through acquisition of convergent mutational sites may leverage conformational adaptability and dynamic couplings between these hotspots to optimize balance between immune evasion and ACE2 affinity.

## Materials and Methods

### Protein Structure Modeling Using AF2 Shallow Subsampling Adaptations

To understand the advantages and limitations of AF2 methods for predicting structural ensembles, we employ (a) MSA subsampling with shallow MSA depth and (b) Random alanine scanning algorithm, which iterates through each amino acid in the native sequence to simulate random alanine substitution mutations. The structural prediction of the Omicron RBD-ACE2 complexes was initially conducted using the AF2 framework within the ColabFold implementation.^80^ This process involved utilizing a range of MSA depths and MSA subsampling techniques. Specifically, we utilized the max_msa field to set two AF2 parameters in the format of *max_seqs:extra_seqs*. These parameters dictate the number of sequences subsampled from the MSA, with *max_seqs* determining the number of sequences passed to the row/column attention track, and extra_seqs determining the number of sequences additionally processed by the main evoformer stack. Lower values in these parameters promote more diverse predictions but may lead to an increased number of misfolded models. The default MSAs are subsampled randomly to obtain shallow MSAs. We set the *max_msa* parameter to 16:32. This parameter is in the format of *max_seqs:extra_seqs* which decides the number of sequences subsampled from the MSA. *Max_seq* determines the number of sequences passed to the row/column attention matrix at the front end of the AF2 architecture, and *extra_seqs* sets the number of extra sequences processed by the Evoformer stack after the attention mechanism. We additionally manipulated the *num_recycles* parameters to produce more diverse outputs. We use *num_recycles*: 12. AF2 makes predictions using 5 models pretrained with different parameters, and consequently with different weights. To generate more data, we set the number of recycles to 12, which produces 14 structures for each model starting from recycle 0 to recycle 12 and generating a final refined structure. Recycling is an iterative refinement process, with each recycled structure getting more precise. Each of the AF2 models generates 14 structures, amounting to 70 structures in total. The MSAs were generated using the MMSeqs2 library using the S sequence as input. We then set the *num_seed* parameter to 1. This parameter quantifies the number of random seeds to iterate through, ranging from *random_seed to random_seed+num_seed*. Increasing the *num_seeds* samples predictions from the uncertainty of the model. Additionally, the dropout parameter was enabled, activating dropout layers in the model during predictions, which further increases variability within the predictions. We also predicted the structure using AF2 with the default and ‘auto’ parameters serving as a baseline structure for prediction analysis.

We also utilized another approach for prediction of multiple conformational states that combined sequence clustering with AF2 and clustered the MSA by sequence distance using DBSCAN method (Density-Based Spatial Clustering of Applications with Noise)^79^. The AF2-cluster approach then runs AF2 predictions using these clusters as input. We employed this approach where MSAs that contained greater than 25% gaps were removed. The rest were clustered using the DBSCAN method, clustering the MSAs by density, and labeling those that were not included as noise and excluding them. This technique generated 13 MSA clusters, which were then used as input for AF2 models using 3 recycles for refinement. As a result, 13 additional structures were predicted using this method. Together, shallow MSA depth and AF2-cluster adaptations generated a total of 83 conformations.

We developed another AF2 adaptation which involves randomized alanine sequence scanning and MSA shallow subsampling. The initial input for the full sequence randomized alanine scanning is the original full native sequence. This technique utilizes an algorithm that iterates through each amino acid in the native sequence and randomly substitutes 5-15% of the residues with alanine, to simulate random alanine substitution mutations. The algorithm substitutes residue a_i_ with alanine at each position i with a probability p_i_ randomly generated between 0.05 and 0.15 for each sequence position. We ran this algorithm nine times on the full native sequence, resulting in nine distinct sequences, each with different frequency and position of alanine mutations. MSAs were constructed for each of these mutated sequences using the alanine-scanned full-length sequences as input for the MMSeqs2 program. The AF2 shallow MSA methodology is then employed on these MSAs to predict protein structures as described previously. A total of 70 predicted structures were generated from 12 recycles per model.

### Statistical and Structural Assessment of AF2-Generated Models

AF2 models underwent ranking based on Local Distance Difference Test (pLDDT) scores, offering a per-residue estimation of prediction confidence ranging from 0 to 100. These scores are calculated by determining the fraction of predicted Cα distances that align with their anticipated intervals. The scores represent the model’s predictions according to the lDDT-Cα metric, which assesses atomic displacements within the predicted model without requiring local superposition.^70,71^ The accuracy of the predicted models was evaluated against the experimental structure using the structural alignment tool TM-align.^81^ This algorithm, specifically developed for sequence-independent protein structure comparison, was utilized to assess and compare the accuracy of protein structure predictions. TM-align iteratively refines the alignment of residues through dynamic programming, aligns and superimposes the structures based on this alignment, and computes the TM-score as a quantitative measure of the overall accuracy of the predicted models. An optimal superposition of the two structures is then built and TM-score is reported as the measure of overall accuracy of prediction for the models. TM-score ranges from 0 to 1, where a value of 1 indicates a perfect match between the predicted model and the reference structure. When TM-score > 0.5 implies that the structures share the same fold. TM-score > 0.5 is often used as a threshold to determine if the predicted model has a fold similar to the reference structure. Several other structural alignment metrics were used including the global distance test total score GDT_TS of similarity between protein structures and implemented in the Local-Global Alignment (LGA) program^82^ and the root mean square deviation (RMSD) superposition of backbone atoms (C, Cα, O, and N) calculated using ProFit (http://www.bioinf.org.uk/software/profit/). This systematic approach provides a quantitative means to assess and rank the structural accuracy of predicted models relative to experimentally determined reference structures.

### All-Atom Molecular Dynamics Simulations

The structures of the Omicron RBD-ACE2 complexes for BA.1 (pdb id 7WBP), BA.2 (pdb id 7XB0), BA.2.75 (pdb id 8ASY), BA.3 (pdb id 7XB1), BA.4/BA.5(pdb id 8AQS) and BQ.1.1 (pdb id 8IF2) are obtained from the Protein Data Bank. In addition, the best AF2 models obtained for each of the Omicron variants were selected as starting structure for MD simulations of RBD-ACE2 complexes. For simulated structures, hydrogen atoms and missing residues were initially added and assigned according to the WHATIF program web interface.^83^ The missing regions are reconstructed and optimized using template-based loop prediction approach ArchPRED.^84^ The side chain rotamers were refined and optimized by SCWRL4 tool.^85^ The protonation states for all the titratable residues of the ACE2 and RBD proteins were predicted at pH 7.0 using Propka 3.1 software and web server.^86,87^ The protein structures were then optimized using atomic-level energy minimization with composite physics and knowledge-based force fields implemented in the 3Drefine method.^88,89^ We considered glycans that were resolved in the structures. NAMD 2.13-multicore-CUDA package^90^ with CHARMM36 force field^91^ was employed to perform 1µs all-atom MD simulations for the Omicron RBD-ACE2 complexes. The structures of the SARS-CoV-2 S-RBD complexes were prepared in Visual Molecular Dynamics (VMD 1.9.3)^92^ and with the CHARMM-GUI web server^9,93^ using the Solutions Builder tool. Hydrogen atoms were modeled onto the structures prior to solvation with TIP3P water molecules^95^ in a periodic box that extended 10 Å beyond any protein atom in the system. To neutralize the biological system before the simulation, Na^+^ and Cl^−^ ions were added in physiological concentrations to achieve charge neutrality, and a salt concentration of 150 mM of NaCl was used to mimic a physiological concentration. All Na^+^ and Cl^−^ ions were placed at least 8 Å away from any protein atoms and from each other. MD simulations are typically performed in an aqueous environment in which the number of ions remains fixed for the duration of the simulation, with a minimally neutralizing ion environment or salt pairs to match the macroscopic salt concentration.^96^

All systems underwent a two-stage minimization protocol. In the first stage, minimization was conducted for 100,000 steps with all hydrogen-containing bonds constrained and the protein atoms fixed. Subsequently, in the second stage, minimization was conducted for 50,000 steps with all protein backbone atoms fixed, followed by an additional 10,000 steps with no fixed atoms. Following minimization, the protein systems underwent equilibration steps by gradually increasing the system temperature in increments of 20 K, ranging from 10 K to 310 K. At each temperature step, a 1 ns equilibration was performed while maintaining a restraint of 10 Kcal mol−1 Å−2 on the protein Cα atoms. After removing restraints on the protein atoms, the system was equilibrated for an additional 10 ns. Long-range, non-bonded van der Waals interactions were computed using an atom-based cutoff of 12 Å, with the switching function beginning at 10 Å and reaching zero at 14 Å. The SHAKE method was used to constrain all the bonds associated with hydrogen atoms. The simulations were run using a leap-frog integrator with a 2 fs integration time step. The ShakeH algorithm in NAMD was applied for the water molecule constraints. The long-range electrostatic interactions were calculated using the particle mesh Ewald method^97^ with a cut-off of 1.0 nm and a fourth-order (cubic) interpolation. The simulations were performed under an NPT ensemble with a Langevin thermostat and a Nosé–Hoover Langevin piston at 310 K and 1 atm. The damping coefficient (gamma) of the Langevin thermostat was 1/ps. In NAMD, the Nosé–Hoover Langevin piston method is a combination of the Nosé–Hoover constant pressure method^98^ and piston fluctuation control implemented using Langevin dynamics.^99,100^ An NPT production simulation was run on equilibrated structures for 1µs keeping the temperature at 310 K and a constant pressure (1 atm).

### Binding Free Energy Computations

We compute the ensemble-averaged binding free energy changes using both conformational ensembles obtained from AF2 predictions and equilibrium samples from simulation trajectories. The binding free energy changes were computed by averaging the results over 1,000 equilibrium samples from MD simulations and 100 AF2-predicted conformations from the obtained conformational clusters. The binding free energies were computed for the Omicron RBD-ACE2 complexes using the Molecular Mechanics/Generalized Born Surface Area (MM-GBSA) approach.^101,102^ We also evaluated the decomposition energy to assess the energy contribution of each amino acid during the binding of RBD to ACE2.^103,104^ The binding free energy for the each RBD–ACE2 complex was obtained using:

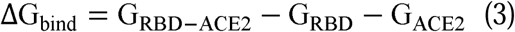

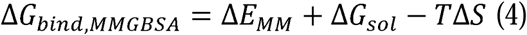

where Δ*E_MM_* is total gas phase energy (sum of Δ*E_internal_*, Δ*E_electrostatic_*, and Δ*Evdw*); Δ*Gsol* is sum of polar (Δ*G_GB_*) and non-polar (Δ*G_SA_*) contributions to solvation.

Here, G_RBD–ACE2_ represent the average over the snapshots of a single trajectory of the MD RBD– ACE2complex (or 100 AF2-predicted conformations), G_RBD_ and G_ACE2_ corresponds to the free energy of RBD and ACE2 protein, respectively. MM-GBSA is employed to predict the binding free energy and decompose the free energy contributions to the binding free energy.^105^ The entropy contribution was not included in the calculations due to the difficulty of accurately calculating entropy for a large protein–protein complex but equally importantly because the entropic differences between variants for estimates of binding affinities are exceedingly small owing to small mutational changes and preservation of the conformational dynamics.

## Results

### Evolutionary and Phylogenetic Analysis of the SARS-CoV-2 Omicron Lineages

We began with a brief analysis of evolutionary differences and divergence among the Omicron variants which are illustrated by the phylogenetic analysis (Figure 2, Supporting Information, Figure S1) using the corresponding clades nomenclature from Nextstrain an open-source project for real time tracking of evolving pathogen populations (https://nextstrain.org/).^106^ Nextstrain provides dynamic and interactive visualizations of the phylogenetic tree of SARS-CoV-2, allowing users to explore the evolutionary relationships between different lineages and variants. Omicron viruses can be divided into two major groups, referred to as PANGO lineages BA.1 and BA.2 or clades 21K and 21L (Figure 2A,B, Supporting Information, Figure S1).

**Figure 2.**
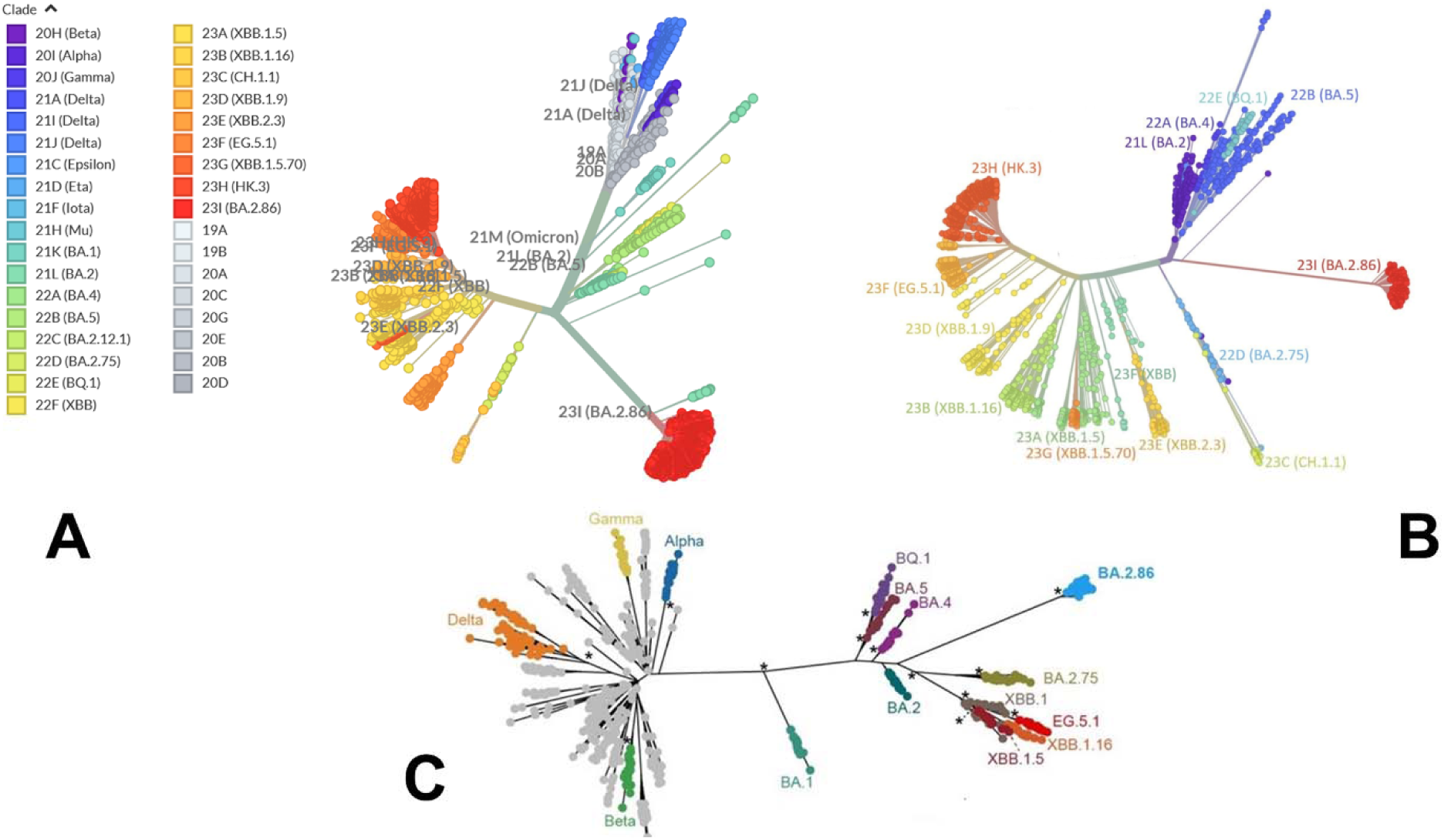
An overview of the phylogenetic analysis and divergence of Omicron variants. The graphs are generated using Nextstrain, an open-source project for real time tracking of evolving pathogen populations (https://nextstrain.org/). (A) The phylogenetic tree of Omicron variants BA.2 (clade 21L), BA.2.75 (clade 22), BA.4 (clade 22A), BA.5 (clade 22B), BA.2.75 (clade 22D), BQ.1 (clade 22E) and XBB lineages including XBB (22F clade) and XBB.1.5 (23A clade). (B) A radial-based phylogenetic tree of Omicron variants highlights evolutionary proximity of BA.2, BA.4, BA.5 and BQ.1.1 variants and more significant evolutionary divergence of BA.2.86 and XBB lineages (C) A phylogenetic tree of Omicron variants relative to Alpha, Beta, Gamma and Delta variants. The evolutionary distance of BA.1, BA.2, BA.4/BA.5 and BQ.1.1 from Alpha, Beta, Gamma and Delta variants is highlighted.

Omicron clades 21K and 21L differ at ∼40 amino acid sites, which is substantial in the context of SARS-CoV-2 evolution while Alpha, Beta and Gamma are as divergent from each other in terms of amino acid changes across the genome as Omicron 21K and 21L are from each other (Figure 2C). BA.3 is not officially recognized by Nextstrain since it is very rare and is labelled as 21M Omicron. In BA.3 21 mutations are shared with all Omicron sub-lineages, Of 21 common mutations, BA.3 shares ten mutations (A67V, H69del, V70del, T95I, V143del, Y144del, Y145del, N211I, L212del, and G446S) from BA.1 and two (S371F and D405N) mutations from BA.2. In other words, of the 33 mutations in the BA.3 lineage spike protein, 31 mutations are common to BA.1. There are no specific mutations for the BA.3 lineage in spike protein. Instead, it is a combination of mutations in BA.1 and BA.2 spike proteins. BA.4 and BA.5 lineages (clades 22A and 22B respectively) have emerged with amino-acid substitutions L452R, F486V, and R493Q reversed in S-protein RBD compared to BA.2 variant (Table 1, Figure 2). Phylogenomic reconstruction indicates that the genomes of BQ.1 (clade 22E) are clustered within the not-monophyletic GSAID Clade 21L, with a close relationship with BA.5 Omicron subvariant (Figure 2B,C). BQ.1 which is a direct descendant of BA.5 has additional spike mutations in some key antigenic sites (K444T and N460K). Its first descendant, BQ.1.1 carries a further additional mutation R346T. In this work, we focus on the analysis of a particular branch of the Omicron tree that includes BA.2, BA.3, BA.4, BA.5 and BQ.1.1 variants (Figure 2, Table 1).

**Table 1.** Mutational landscape of the Omicron variants.

| Omicron Variant | Mutational landscape |
| --- | --- |
| BA.1 | T95I, G339D, S371L, S373P, S375F, K417N, N440K, G446S, S477N, T478K, E484A, Q493R, G496S, Q498R, N501Y, Y505H, T547K, D614G, H655Y, N679K, P681H, N764K, D796Y, N856K, Q954H, N969K, L981F |
| BA.2 | T19I, G142D, V213G, G339D, S371F, S373P, S375F, T376A, D405N, R408S, K417N, N440K, S477N, T478K, E484A, Q493R, Q498R, N501Y, Y505H, D614G, H655Y, N679K, P681H, N764K, D796Y, Q954H, N969K |
| BA.2.75 | T19I, G142D, K147E, W152R, F157L, I210V, V213G, G257S, G339H, S371F, S373P, S375F, T376A, D405N, R408S, K417N, N440K, G446N, N460K, S477N, T478K, E484A, Q498R, N501Y, Y505H, D614G, H655Y, N679K, P681H, N764K, D796Y, Q954H, N969K |
| BA.3 | G142D, G339D, S371F, S373P, S375F, D405N, K417N, N440K, G446, S477N, T478K, E484A, Q493R, Q498R, N501Y, Y505H, D614G, H655Y, N679K, P681H, N764K, D796Y, Q954H, N969K |
| BA.4 | T19I, G142D, V213G, G339D, S371F, S373P, S375F, T376A, D405N, R408S, K417N, N440K, L452R, S477N, T478K, E484A, F486V, R493Q reversal, Q498R, N501Y, Y505H, D614G, H655Y, N679K, P681H, N764K, D796Y, Q954H, N969K |
| BA.5 | T19I, LPPA24-27S, Del 69-70, G142D, V213G, G339D, S371F, S373P, S375F, T376A, D405N, R408S, K417N, N440K, L452R, S477N, T478K, E484A, F486V, R493Q reversal, Q498R, N501Y, Y505H, D614G, H655Y, N679K, P681H, N764K, D796Y, Q954H, N969K |
| BQ.1.1 | T19I, LPPA24-27S, H69del, V70del, V213G, G142D, G339D, R346T, S371F, S373P, S375F, T376A, D405N, R408S, K417N, N440K, K444T, L452R, N460K, S477N, T478K, E484A, F486V, R493Q reversal, Q498R, N501Y, Y505H, D614G, H655Y, N679K, P681H, N764K, D796Y, Q954H, N969K |

### AF2 Atomistic Modeling and Structural Prediction of the Omicron RBD-ACE2 Complexes

Despite considerable mutational differences between newly emerged Omicron variants, structural analysis of the RBD complexes with ACE2 for these variants revealed similar RBD conformations and the same binding mode of interactions. We embarked on a systematic comparative analysis of conformational dynamics and energetics of the Omicron RBD-ACE2 complexes using a panel of evolutionary proximal BA.1, BA.2, BA.2.75, BA.3, BA.4/BA.5 and BQ.1.1 variants. To facilitate this comparative analysis, we explored both AF2-based adaptations for modeling of structural ensembles and MD simulations to characterize conformational landscapes and functional conformational states of the Omicron spike proteins. We began with baseline AF2 structural predictions of the Omicron RBD-ACE2 complexes using AF2 within the ColabFold^80^ and best five models for each system (Figure 3). The confidence of the AF2 predictions for RBD-ACE2 complexes is analyzed using residue-based pLDDT scores (Figure 3).

**Figure 3.**
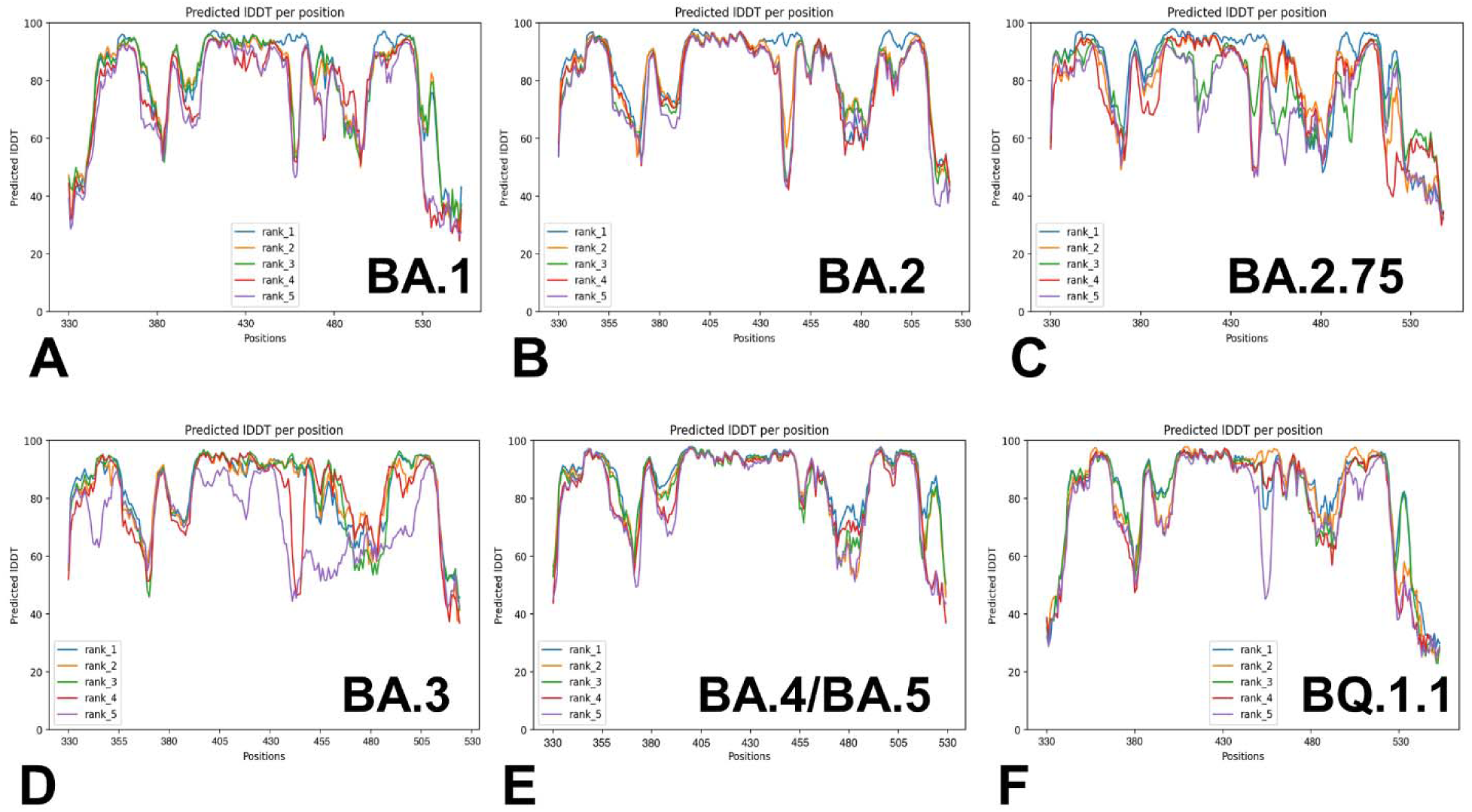
The AF2 analysis of predictions for the Omicron RBD-ACE2 The pLDDT per residue for the top five models obtained from AF2 predictions of the BA.1 RBD-ACE2 complex (A), BA.2 RBD-ACE2 complex (B), BA.2.75 RBD-ACE2 complex (C), BA.3 RBD-ACE2 complex (D), BA.4/BA.5 RBD-ACE2 complex (E) and BQ.1.1 RBD-ACE2 complex (F). pLDDT between 70 and 90 indicate a high accuracy, where the prediction of the main chain of the protein is reliable. pLDDT values above 90 indicate extremely high accuracy, equivalent to structures determined by experiments. pLDDT values between 50 to 70 indicate a lower accuracy, but it is likely that the predictions of individual secondary structures are correct.

The results showed a significant convergence among five independent AF2 runs for each complex. However, this pattern is particularly evident for BA.2 (Figure 3B), BA.4/B.5 (Figure 3E) and BQ.1.1 complexes (Figure 3F). We noticed also appreciable divergence between the pLDDT values of the top five models for these regions. These preliminary results suggested that AF2-generated top structural models can accurately reproduce the RBD fold and stability of the RBD core regions. Moreover, we argue that the lower pLDDT values obtained for the RBD loops and particularly RBM resides reflected the intrinsically flexible and adaptable nature of these regions rather than a reduced prediction quality of the AF2 pipeline. Structural alignment of the best five AF2 models with the experimental structures for the Omicron RBD-ACE2 complexes yielded the RMSD values of ∼0.5-0.8 Å (Figure 4), suggesting high prediction accuracy for all systems. In addition, we highlighted the pLDDT confidence values and RMSDs for the best model, showing that pLDDT scores were above 80.0 -82.0 and the RMSD values were below 0.5 Å (Figure 4). A comparison of the top predicted models with the experimental structures demonstrated the high accuracy of predictions and highlighted only small deviations in the intrinsically flexible RBM loop (residues 475-487) and in peripheral flexible region (residues 520-527) (Figure 4). Importantly, the AF2 results for all Omicron RBD-ACE2 complexes showed that selection of best models based on the highest pLDDT scores can yield functionally relevant mobility in the RBD loop 444-452 and RBM tip 475-487 that harbor important mutational sites across all Omicron variants. Although there are some minor differences in the amplitude of the RBM tip fluctuations among variants, the predicted spectrum of RBM conformations displayed an ordered “hook-like” folded RBM tip which is a preferable state in the experimental structures. Previous structural and computational studies suggested that a hook-like folded RBD tip (the “Hook” state) may interconvert during equilibrium with less dominant and highly dynamic “disordered” state in which the RBD tip cycles between a variety of conformations.^119^ Importantly, the best AF2 models preserved the hook-like folded RBM conformations only revealing modest lateral displacements (Figure 4).

**Figure 4.**
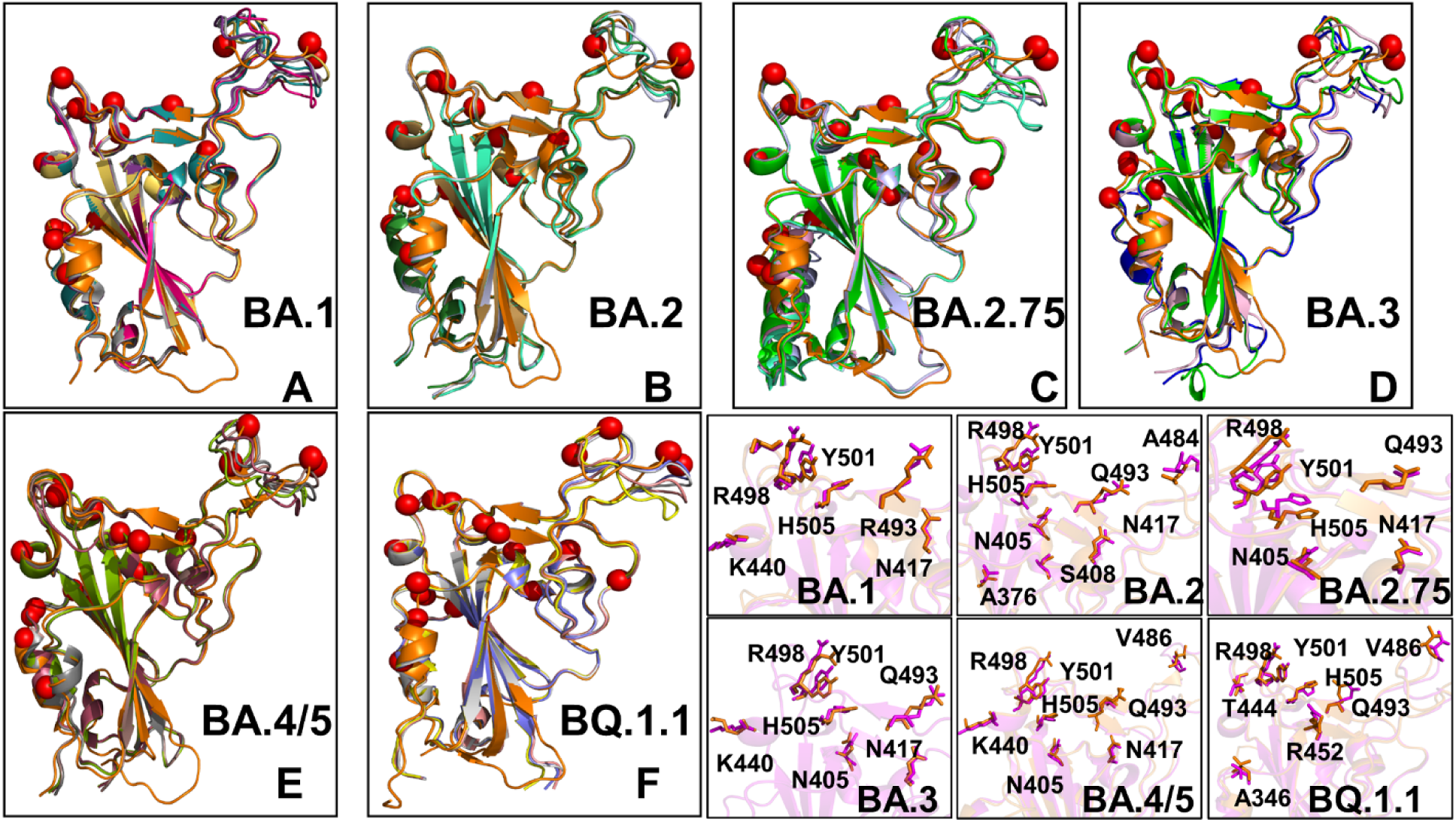
Structural alignment of the AF2-predicted top five models with the experimental structure for the Omicron RBD-ACE2 complexes. (A) Structural alignment of the AF2-predicted top five BA.1 conformations with highest pLDDT values and the experimental structure of the BA.1 RBD-ACE2 complex (in orange ribbons), pdb id 7WBP. (B) Structural alignment of the AF2-predicted top five BA.2 conformations and the experimental structure of the BA.2 RBD-ACE2 complex (in orange ribbons), pdb id 7XB0. (C) Structural alignment of the AF2-predicted top five BA.2.75 conformations and the experimental structure of the BA.2.75 RBD-ACE2 complex (in orange ribbons), pdb id 8ASY. (D) Structural alignment of the AF2-predicted top five BA.3 conformations and the experimental structure of the BA.3 RBD-ACE2 complex (in orange ribbons), pdb id 7XB1. (E) Structural alignment of the AF2-predicted top five BA.4/BA.5 conformations and the experimental structure of the BA.4/5 RBD-ACE2 complex (in orange ribbons), pdb id 8AQS. (F) Structural alignment of the AF2-predicted top five BQ.1.1 conformations with high pLDDT values and the experimental structure of the BQ.1.1 RBD-ACE2 complex (in orange ribbons), pdb id 8IF2. The RBD conformations are shown in ribbons and the positions of the Omicron RBD mutational sites for each of the respective variants are shown in red spheres. (Bottom right panel). The closeups of the predicted side-chains (shown in magenta sticks) and the experimental conformations ( in orange sticks) for the Omicron mutational sites in BA.1, BA.2, BA.2.75, BA.3, BA.4/BA.5 and BQ.1.1 variants.

It is worth noting that dynamic changes in the RBM tip may affect conformations and interacting positions for key mutational sites for making favorable interactions with ACE2 for mutational sites E484A and F486 (or F486V in BA.4/BA.5 and BQ.1.1). In the structural context of the full length trimer, conformational mobility of the RBM tip may promote the increased population of the RBD-up states and modulate binding interactions with ACE2 and antibodies.^119^ Similar functional variations of the RBM residues were also observed in MD simulations showing that the RBM loop has an inherent conformational flexibility that is not observed in the static structures and that ACE2 and antibody binding to this region may elicit specific distribution of conformations as compared to the unbound RBD form.^107^ The predicted plasticity of the RBM tip in the top AF2 models is functionally relevant and consistent with the hydrogen/deuterium-exchange mass spectrometry (HDX-MS) studies of spike flexibility^108–110^ particularly showing bimodal isotopic distributions between two distinct interconverting populations of ACE2-bound structurally stable and more flexible RBM conformations.^109^

### AF2 Adaptations with Shallow MSA and Randomized Sequence Scanning Enable Predictions of Protein Conformational Ensembles for the Omicron RBD-ACE2 Complexes

We used AF2 adaptation with varied MSA depth to predict structural ensembles of the RBD-ACE2 complexes. In this analysis, it is assumed that the experimental structure would be the best ( or among best) ranked models within the ensemble based on the pLDDT metric assessment. The density distribution of the pLDDT scores for the ensemble of the AF2 models revealed single pronounced cluster peaks at pLDDT ∼ 80-90 for BA.2 (Figure 5B), BA.4/BA.5 (Figure 5E) and BQ.1.1 variants (Figure 5F). This implies that the largest fraction of the ensemble samples a narrow basin around the native state. Interestingly, for BA.1 (Figure 5A), BA.2.75 (Figure 5C) and BA.3 (Figure 5D) the distributions are broader featuring peaks pLDDT ∼65-75 and pLDDT∼ 80-85 indicating presence of several conformational clusters within the native RBD fold with the potentially increased variability of the RBD flexible regions.

**Figure 5.**
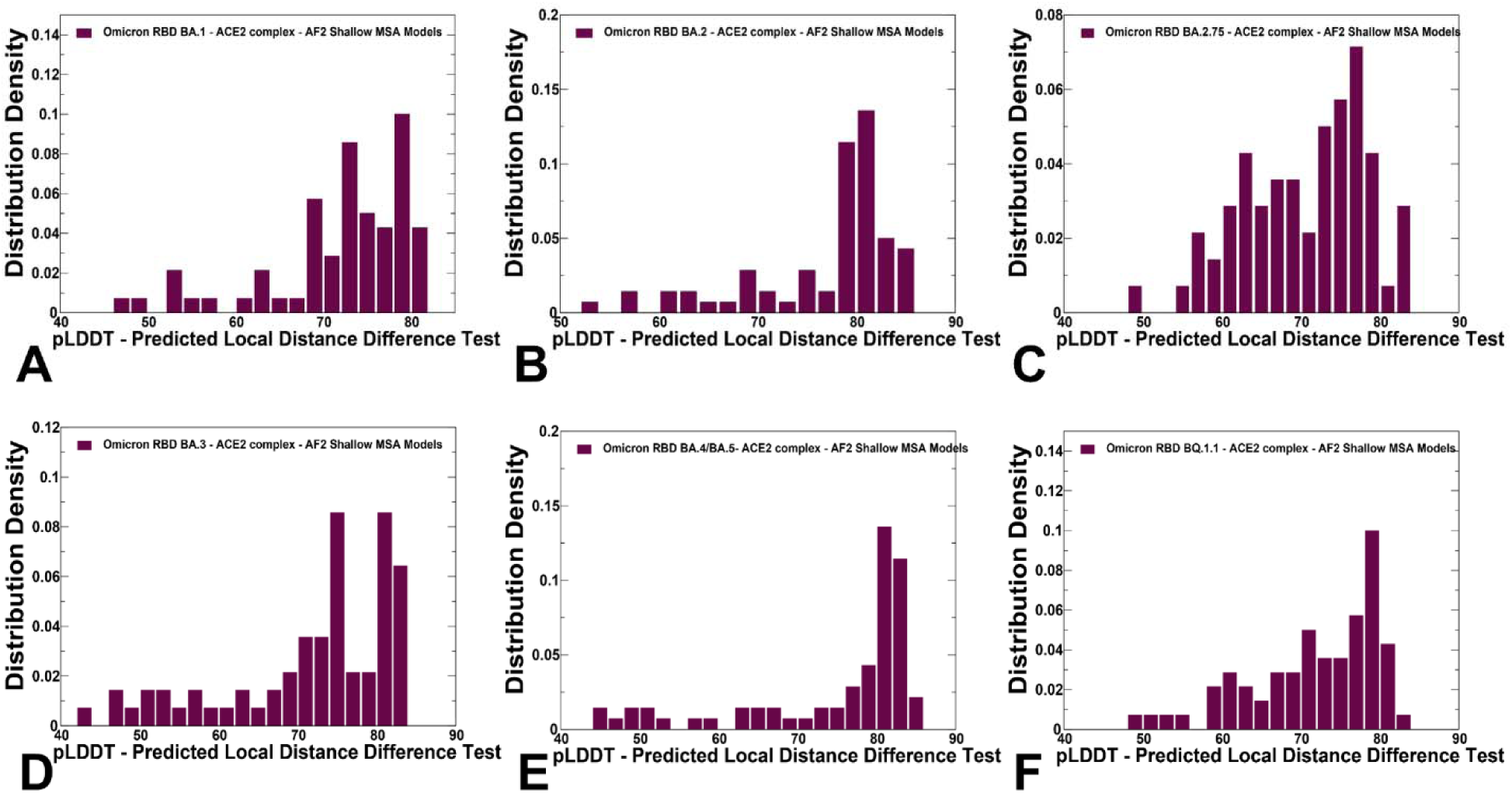
The distributions of the pLDDT metric for the RBD-ACE2 conformational ensembles obtained from AF2-MSA shallow depth predictions. The density distribution of the pLDDT values for structural ensembles are shown for BA.1 (A) BA.2 (B), BA.2.75 (C), BA.3 (D), BA.4/BA.5 (E) and BQ.1.1 (F). The density distributions are depicted as maroon-colored filled bars.

Notably, for all distributions, there is a sharp decay of the density at the lower pLDDT values ∼50-70 reflecting minor population of highly flexible RBD conformations with the increased mobility of the loop regions and fluctuations of the RBD core (Figure 5). AF2 predictions with pLDDT values ∼70-90 are typically associated with high confidence, the regions with pLDDT values ∼ 50-70 have lower confidence, while pLDDT < 50 may be a strong predictor of disorder. According to our results, there was no measurable disorder in the predicted conformations and yet emergence of the native conformations with an appreciable plasticity in the flexible sites.

Using structural similarity metrics TM-score and RMSD the prediction accuracy of AF2-MSA depth models was further evaluated. The density distributions of TM-scores showed a considerable similarity between the predicted conformations and the experimental structures with the major peaks corresponding to TM-score ∼0.9 (Figure 6). For BA.2, BA.4/BA.5 and BQ.1.1 variants the TM score distribution revealed clusters at TM score ∼0.9-0.95 thereby showing a strong preference for the native structure. Broader distributions were observed for BA.1, BA.2.75 and BA.3 (Figure 6A,C,D) showing presence of several conformational clusters with TM score spanning range between 0.7 and 0.95 values.

**Figure 6.**
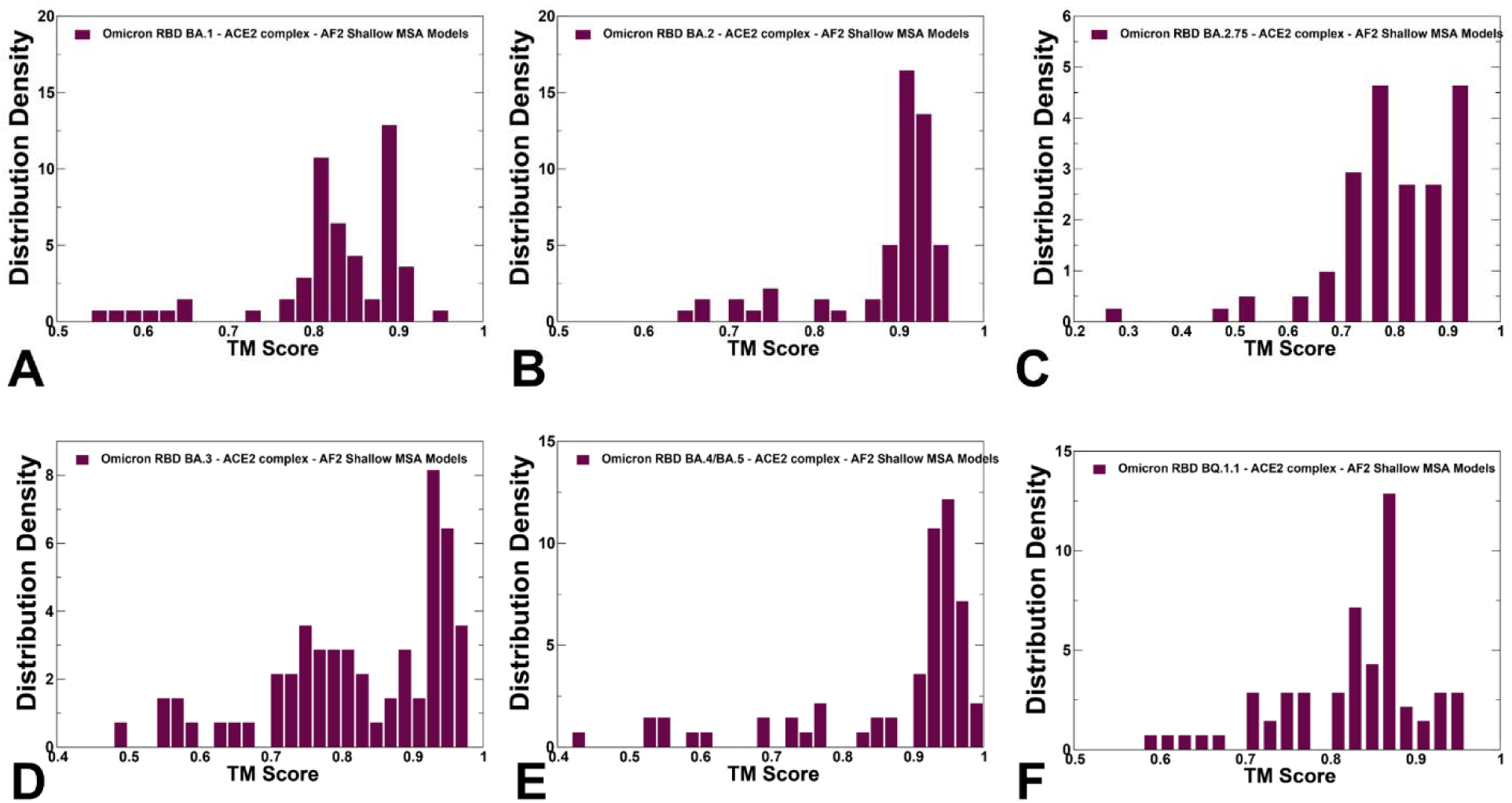
The density distribution of TM-score measuring structural similarity of the AF2-MSA shallow depth predicted ensembles with respect to the experimental structures. The density distribution of the of TM-scores for BA.1 (A) BA.2 (B), BA.2.75 (C), BA.3 (D), BA.4/BA.5 (E) and BQ.1.1 (F). The density distributions are shown in maroon-colored filled bars.

A consistent picture emerged from the density distribution of the RMSDs for the predicted ensemble from the experimental structures (Figure 7) showing dominant cluster peaks at RMSD∼ 0.6-0.8 Å and reduced density at RMSDs ∼1.0-1.2 Å. Overall, for all variants, we observed that the predicted AF2 conformations sample the native basin around the experimental structures. We also examined the relationship between pLDDT confidence estimates and RMSD values of the predicted conformations from the experimental structures (Figure 8). A strong correlation was obtained for all systems showing that the higher pLDDT value of the predicted conformation the lower the RMSD value for this model from the experimental structure (Figure 8). The Pearson correlation coefficients were significant for all variants. Another interesting feature of the scatter profiles is a clear evidence of highly populated clusters of native conformations with pLDDT values ∼75-85 and RMSD < 0.7 Å (Figure 8).

**Figure 7.**
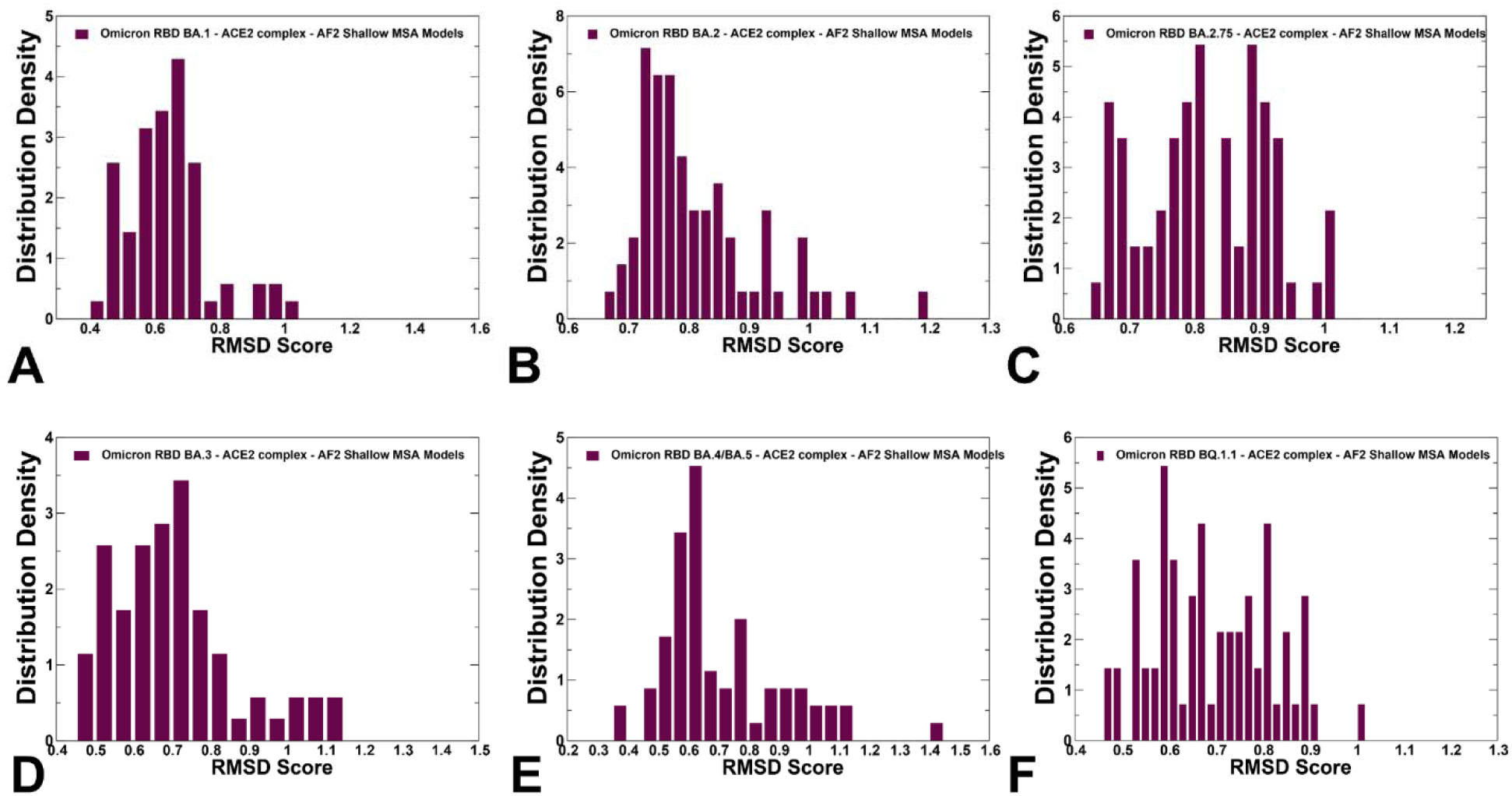
The density distribution of the RMSD scores measuring structural similarity of the AF2-MSA shallow depth predicted RBD conformational ensembles with respect to the experimental structures. The density distribution of the of the RMSD scores for structural ensembles are shown for BA.1 (A) BA.2 (B), BA.2.75 (C), BA.3 (D), BA.4/BA.5 (E) and BQ.1.1 (F). The density distributions are depicted as maroon-colored filled bars.

**Figure 8.**
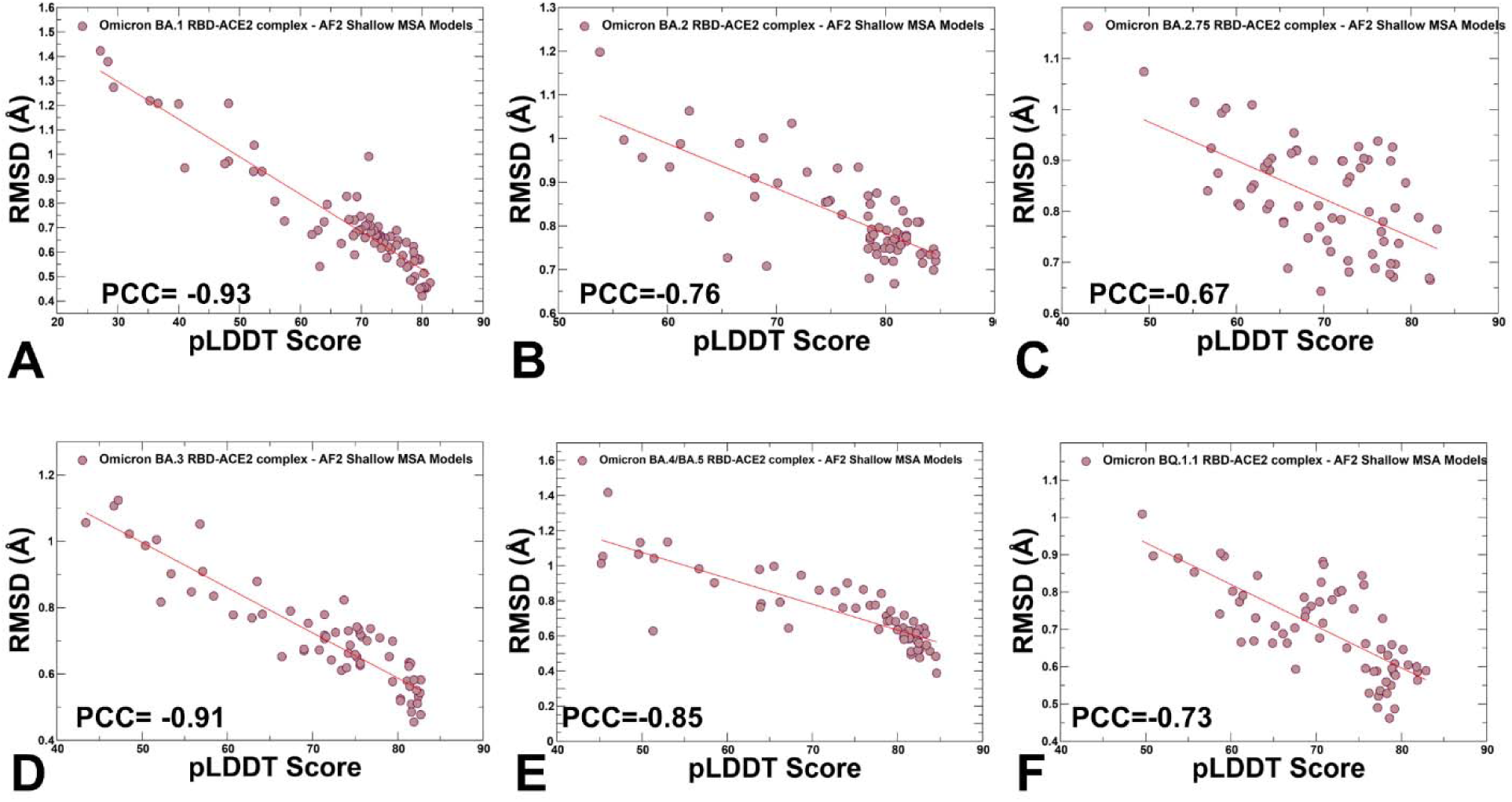
The scatter plots between the pLDDT scores and RMSD scores measuring confidence and structural similarity of the AF2-MSA shallow depth predicted RBD conformational ensembles with respect to the experimental structures. The scatter plot distributions are shown for structural ensembles obtained for BA.1 (A) BA,2 (B), BA.2.75 (C), BA.3 (D), BA.4/BA.5 (E) and BQ.1.1 (F). The density distributions are depicted as brown-colored filled circles. The Pearson correlation coefficients (PCC) values between the pLDDT and RMSD values are shown in respective panels.

As pLDDT values decrease below this threshold, more dispersion of RMSD values and greater variability can be observed for conformations with RMSDs > 1.0 Å (Figure 8). Hence, the highest pLDDT-scoring models typically correspond to the native conformations which implies that selection of protein conformations based on the pLDDT values can enable identification of functionally relevant conformations. Although the AF2 models generated using shallow MSA approach produced clusters of functional conformations near the native state, the diversity of the generated ensembles is still relatively limited due to strong bias of the AF2 models towards the native state. To explore this issue, we first employed the AF2-cluster approach that generates a MSA with ColabFold^79^, following by clustering MSA sequences with DBSCAN, and running AF2 predictions for each cluster (Supporting Information, Figure S2). This AF adaptation was designed to capture conformational variability by altering input MSAs, and our results indicate the increased diversity of predicted conformations. By inspecting the pLDDT, TM-score, and RMSD values of the generated conformational clusters for BA.2 and BQ.1.1 variants (Supporting Information, Figure S2), we observed pLDDT ∼ 60-70 for some clusters but significantly lowered pLDDT ∼ 40-60 indicating reduced confidence level due to elevated flexibility of the RBD conformations. Similarly, while most AF2-cluster conformations yielded reasonable TM-score values > 0.65-0.7 for native-like RBD states, several predicted conformations with TM-scores ∼-0.4-0.6 highlighted the elevated variability across all RBD regions (Supporting Information, Figure S2). The RMSD values for the cluster conformations reflected the increased variability consistent with the trend revealed by the pLDDT analysis (Supporting Information, Figure S2). Overall, the AF-cluster analysis suggested that generated RBD conformations may significantly differ, as in addition to functionally relevant RBD cluster conformations, this approach can produce conformations with the amplified flexibility and even generate partially disordered states.

To better characterize functionally relevant conformational states and structural ensembles using the AF2 framework, we proposed randomized alanine scanning adaptation of the AF2 methodology in which the algorithm operates on sequences and iterates through each amino acid in the native sequence to randomly substitute 5-15% of the residues with alanine, thus emulating random alanine mutagenesis. A comparison of the pLDDT profiles highlighted a clear shift and broadening of the distribution using randomized scanning AF2 adaptation, yielding pLDDT values ∼60-80 (Supporting Information, Figure S3). Accordingly, the protein conformational ensembles generated by this AF2 approach revealed a more extensive heterogeneity of predicted states, yielding RBD states with RMSDs ∼ 1.0-1.7 Å from the experimental structures (Supporting Information, Figure S4). A comparison of the RMSD distributions for protein conformations produced by AF2 with shallow MSA and randomized sequence scanning approaches pointed to their complementarity enabling to expand the AF2-based ensemble. The scatter plots between pLDDT and RMSD values for this expanded ensemble retained a strong correlation (Supporting Information, Figure S5) by revealing several clusters of conformations associated with the higher RMSD ∼ 1.0-1.7 Å and lower pLDDT values ∼50-70. Hence, by combining the AF2-genersted ensembles can expand the scope of sampled conformational states. Moreover, the results showed that the inverse correlation between pLDDT and RMSD values can be mostly preserved in the extended ensemble (Supporting Information, Figure S5), confirming that the pLDDT metric can serve as a robust indicator of the functional conformational ensembles. Based on the results of this analysis we argue that by selecting AF2-generated conformations produced by both AF2 schemes with high confidence pLDDT values ∼70-90 the predicted pool of functional RBD conformations can be judiciously expanded.

Structural alignment of the AF2 conformational ensembles illustrated the predicted patterns of RBD mobility in which the RBD core and most of the loops largely remain in their native positions, while most of variability is observed in the RBD loop 457-475, RBM loop 475-487 and peripheral flexible region (residues 520-527) (Figure 9). This flexible region is immediately next to the C-terminal domain CTD1 (residues 529-591) in the full length S trimer that functions as a structural relay between RBD and S2 regions that can arguably sense the functional movements in the S1 and S2 subunits. The flexible region 520-527 in the trimer is a part of the RBD-CTD1 hinge connecting the C-terminal of the RBD with CTD1 (C-connector). Together with the N-connector located near the N-terminal of the RBD, these flexible regions can modulate RBD openings during functional changes between the closed and open RBD states. Instructively, we found that the conformational ensembles for BA.2 (Figure 9B), BA./BA.5 (Figure 9E) and BQ.1.1 complexes (Figure 9F) displayed moderate variations of the RBD loops and were mostly confined to the native state. At the same time, more heterogeneous conformational ensembles were obtained for BA.1 (Figure 9A) and BA.3 variants (Figure 9D). Interestingly, one of the most dynamic regions of the RBD (loop 457-475) is immediately adjacent to residues L455 and F456 that emerged among convergent mutational hotspots in the XBB variants. ^111–113^ Based on these findings, we argue that the AF2-predicted conformational plasticity in the RBM region may be functionally significant. Importantly, pLDDT assessment of the AF2 models can be used to quantify the variability and extent of dynamic changes in the flexible RBD regions. Although AF2 predictions do not necessarily represent the equilibrium ensemble of conformations, our results suggested that the generated conformational states may represent functionally significant representatives of the equilibrium ensemble of RBD conformations. In general, these results showed that AF2-generated conformational ensembles can accurately reproduce the experimental structures and capture conformational details of the RBD fold and variant-specific functional adjustments of the RBM loop impacting the exposure for mutational positions in this region.

**Figure 9.**
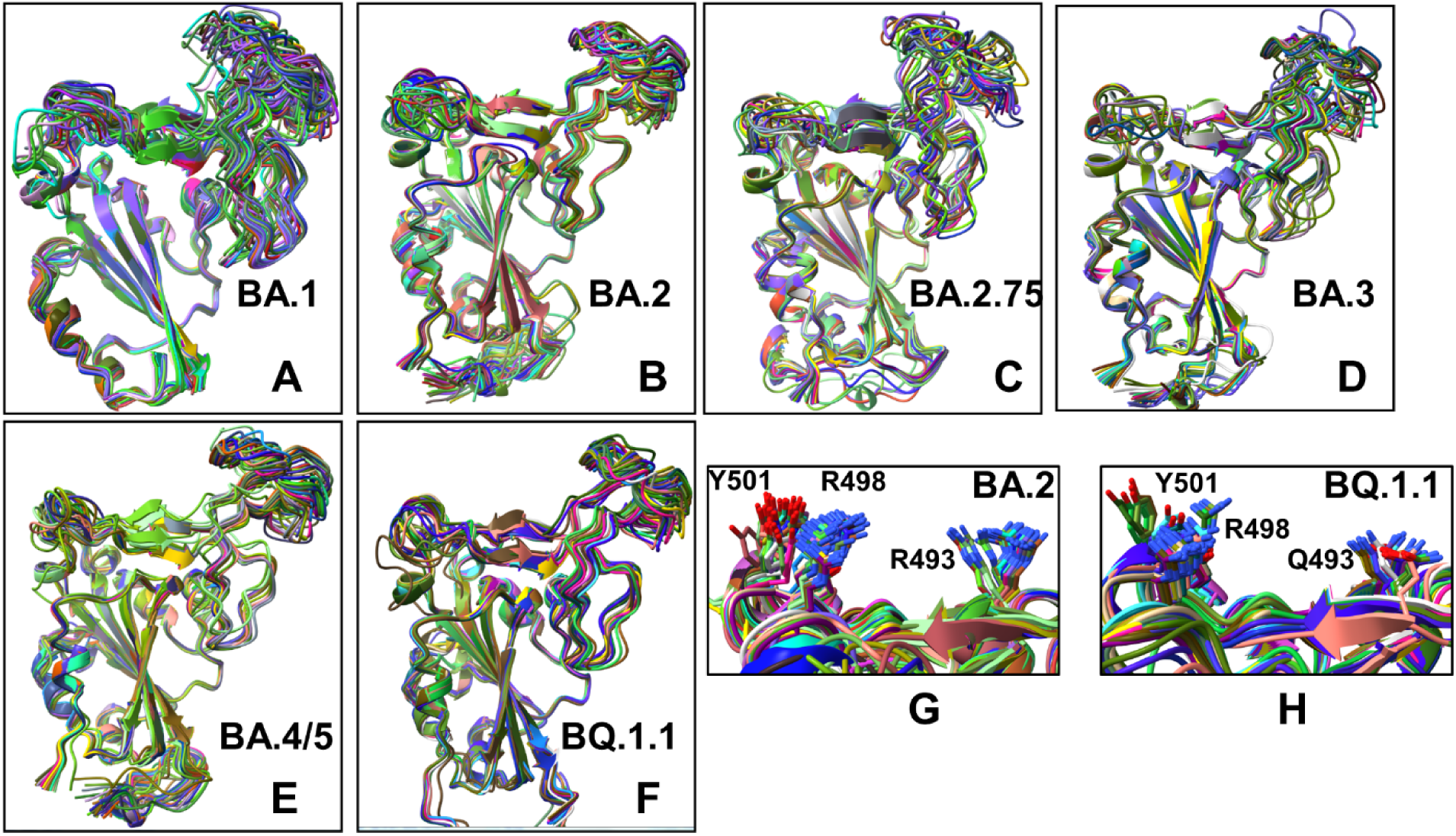
Structural alignment of the AF2-generated conformational ensembles obtained by AF2 shallow MSA approach and AF2 adaptation with randomized alanine sequence scanning. Structural alignment of the AF2-predicted ensemble of RBD conformations with the respective experimental structures (shown in orange ribbons) for the BA.1 RBD-ACE2 complex (A), BA.2 RBD-ACE2 complex (B), BA.2.75 RBD-ACE2 complex (C), BA.3 RBD-ACE2 complex (D), BA.4/BA.5 RBD-ACE2 complex (E), and BQ.1.1 RBD-ACE2 complex (F). The RBD conformations are shown in ribbons. (G) A closeup of the side-chain conformations in the predicted ensemble of BA.2 RBD-ACE2 complex for the key Omicron mutational sites R493, R498 and Y501. The residue conformations are shown in sticks. (H) A closeup of the side-chain conformations in the predicted ensemble of BQ.1..1 RBD-ACE2 complex for the key Omicron mutational sites R493Q, R498 and Y501. The residue conformations are shown in sticks.

### Atomistic MD Simulations of the XBB RBD-ACE2 ACE2 Complexes

To characterize conformational landscapes and dynamic signatures of the Omicron RBD-ACE2 variants we also conducted microsecond MD simulations (Figure 10). Conformational dynamics profiles obtained from MD simulations were similar and revealed several important trends. The RMSF profiles showed local minima regions corresponding to the structured five-stranded antiparallel β-sheet core region that functions as a stable core scaffold (residues 350-360, 375-380, 394-403) and the interfacial RBD positions involved in the contacts with the ACE2 receptor (Figure 10A,B). The RMSF profiles also displayed common RMSF peaks corresponding to the flexible RBD regions including residues 360-373, residues 380-396 as well as RBM tip residues 475-487. As expected, the conformational dynamics profiles also showed the increased RMSF values in the flexible N-terminal and C-terminal ends of the RBD structure, but fluctuations in these regions were moderate as compared to more elevated mobility of C-terminal RBD region (residues 515-527) in the AF2 predictions (Figure 10A,B). The RBD core regions (residues 390-420, 430-450) exhibited small fluctuations, particularly in the BA.2 and BA.2.75 variants (Figure 10A) suggesting the increased RBD stability for these variants which may be a relevant contributing factor to the experimentally observed stronger ACE2 binding. In addition, the RMSF profiles are characterized by several local minima corresponding to the ACE2 interfacial sites (residues 485-505). Consistent with AF2 predictions, the MD profiles displayed larger displacements in the flexible RBD regions (residues 355-375, 381-394, 444-452, 455-471, 475-487,515-527) for BA.3 and BA.4/BA.5 complexes (Figure 10A,B). Despite significant similarities of the RMSF profiles, we found several notable differences that are particularly important in the context of the AF2 predictions.

**Figure 10.**
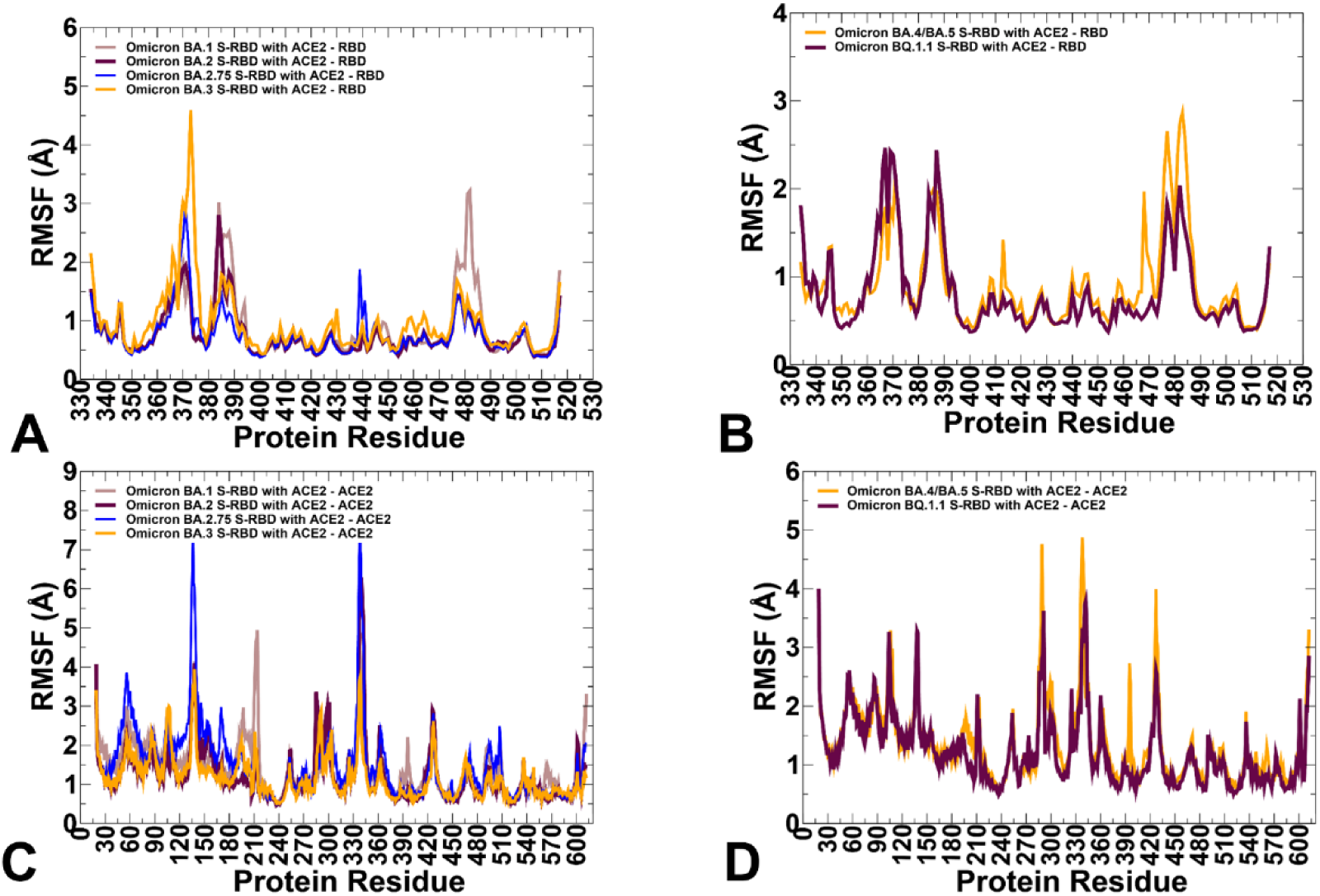
Conformational dynamics profiles obtained from all-atom MD simulations of the Omicron RBD complexes with ACE2. (A) The RMSF profiles for the RBD residues obtained from MD simulations of the BA.1 RBD-ACE2 complex, pdb id 7WBP ( in light brown lines), the BA.2 RBD-ACE2 complex, pdb id 7XB0 (in maroon lines), BA.2.75 RBD-ACE2 complex, pdb id 8ASY (in blue lines) and BA.3 RB-ACE2 complex, pdb id 7XB1 (in orange lines). (B) The RMSF profiles for the RBD residues obtained from MD simulations of the BA.4/BA.5 RBD-ACE2 complex, pdb id 8AQS (in orange lines) and the BQ.1.1 RBD-ACE2 complex, pdb id 8IF2 (in maroon lines). (C) The RMSF profiles for the ACE2 residues obtained from MD simulations of the BA.1 RBD-ACE2 complex ( in light brown lines), the BA.2 RBD-ACE2 complex (in maroon lines), BA.2.75 RBD-ACE2 complex (in blue lines) and BA.3 RB-ACE2 complex (in orange lines). (D) The RMSF profiles for the ACE2 residues obtained from MD simulations of the BA.4/BA.5 RBD-ACE2 complex, pdb id 8AQS (in orange lines) and the BQ.1.1 RBD-ACE2 complex, pdb id 8IF2 (in maroon lines).

The dynamics profiles showed the markedly larger fluctuations of the RBM tip region (residues 475-487) in the BA.1 variant and smallest fluctuations in this region for BA.2.75 variant (Figure 10A). We also observed smaller fluctuations in BA.2.75 for the hairpin loop (residues 373– 380).These findings are consistent with the experimental studies showing that the reversion of R493Q and local conformational alterations in the hairpin loop 373–380 can result in the overall improved stability for the RBD and the RBM region.^35^ It is worth noting that the AF2-generated ensembles for BA.2.75 variant showed more conformational plasticity as compared to MD simulations and produced high confidence RBD states with lateral changes in the RBB tip. MD simulations also highlighted the moderately increased flexibility of the BA.4/BA.5 RBD as compared to more stable BQ.1.1 variant (Figure 10B). These observations are in line with structural analysis showing that F486V in the BA.4/5 S RBD may decrease the hydrophobic interaction with ACE2 and promote greater mobility of the RBM region, while Q493 forms a hydrogen bond with the ACE2 residue H34 can compensate for the loss of binding.^38,39^ Consistent with this analysis, conformational dynamics profile of the BA.4/BA.5 RBD-ACE2 complex revealed markedly increased flexibility of the RBM region where the RBM tip residues, including F486V position can undergo RMSF changes ∼ 2.5-3.0 Å (Figure 10B). Although the distribution of rigid and flexible RBD regions is preserved and shared across all RBD complexes, the extent of stability and mobility may be modulated to indue the increased rigidity of several stable RBD regions in BA.2, BA.2.75 and BQ.1.1 variants. The conformational dynamics profile of the ACE2 receptor showed a similar and strong stabilization of the interfacial helices on ACE2, indicating that dynamics signatures of ACE2 are conserved across all Omicron RBD complexes (Figure 10C,D). We observed the increased structural stability of ACE2 residues in BA.2 (Figure 10C) and BQ1.1 (Figure 10D). While the stability of the ACE2 interfacial regions (residues 350–395) is stronger in the BA.2 and BA.2.75 complexes, MD simulations revealed larger fluctuations in the flexible ACE2 regions for these variants which may reflect long-range communications between the binding interface and peripheral regions in the ACE2.

In general, a comparison of AF2 ensemble predictions and MD simulations suggested that MD profiles can be more sensitive to mutational changes and depict subtle changes in the conformational stability between variants. The range of thermal fluctuations in the MD-generated conformational ensembles is more limited than in the AF2 ensembles and confined to the close proximity of the native state. In some contrast, AF2-generated conformational ensembles may produce a broader pool of the energetically favorable and diverse RBD conformations, enabling a better representation of functional RBM displacements and movements of the flexible RBD loops during binding with the ACE2 receptor. Despite the increased mobility, the RBM tip maintains a stable folded “hook” conformation that is similar to the cryo-EM conformations.

These results suggested complementarities and potential synergies between AF2 predictions of protein conformational ensembles and MD simulations showing that integrating information from both methods can potentially yield a more adequate characterization of the conformational landscapes for the Omicron RBD-ACE2 complexes. Noteworthy, the nature of the AF2 adaptations for probing structural ensembles rather than predicting a single structure is based on different manipulations of the MSAs which drive diversity of conformational predictions without necessarily rigorously considering the underlying thermodynamic distributions. Nonetheless, by evaluating the quality of the generated conformations using pLDDT metric and considering conformations with high confidence pLDDT ∼70-90 these tools can greatly expand coverage of the accessible conformational landscape and identify functionally relevant clusters of states. Ideally, by launching MD runs from a diverse set of the AF2-predicted conformational pool, simulations can reweight structural distributions and provide a physically rigorous assessment of most probable functional conformations. Hence, combining AF2 predictions with MD simulations may enable a more robust and accurate characterization of dynamic binding mechanisms.

### MM-GBSA Analysis of the Binding Affinity Computations for the RBD-ACE2 Complexes

Using the conformational ensembles obtained from AF2 predictions and MD simulations, we computed the binding free energies for the RBD-ACE2 complexes using the MM-GBSA method. ^101–104^ The results of MM-GBSA computations using MD-based equilibrium ensemble showed a good agreement with the experimental SPR-measured binding affinities for the RBD-ACE2 complexes (Table 2). Interestingly, the decomposition of binding free energy terms showed relatively similar contributions of the van der Waals and electrostatic interactions in the BA.2 and BA.2.75 RBD-ACE2 complexes that yielded the best binding free energies (Table 2).

**Table 2.** MM-GBSA Binding Energies for the RBD-ACE2 Complexes Using MD Ensembles.

| **System** | $E_{\text{vdW}}$ | $E_{\text{elec}}$ | $E_{\text{GB}}$ | $E_{\text{SA}}$ | $\Delta G_{\text{bind}}$<br>(kcal/mol) | Exp.binding |
| --- | --- | --- | --- | --- | --- | --- |
| BA.1 RBD-ACE2 | -102.42 | -1420.36 | 1453.71 | -13.35 | -82.42 | 19.5 nM |
| BA.2 RBD-ACE2 | -113.81 | -1474.06 | 1514.1 | -14.82 | -88.58 | 10.0 nM |
| BA.2.75 RBD-ACE2 | -113.47 | -1475.26 | 1512.98 | -14.75 | -90.05 | 8.21 nM |
| BA.3 RBD-ACE2 | -102.97 | -1453.82 | 1492.93 | -13.79 | -83.94 | 22 nM |
| BA.4/BA.5 RBD-ACE2 | -84.23 | -1337.37 | 1344.24 | -10.95 | -88.31 | 14.37 nM |
| BQ.1.1 RBD-ACE2 | -95.51 | -1231.18 | 1251.38 | -12.74 | -88.06 | 14.70 nM |

These energetic contributions are also the most favorable for BA.2 and BA.2.75 complexes. The analysis revealed that for BA.1, BA.2, BA.2.75 and BA.3 variants the increased electrostatic contributions are positively correlated with the enhanced ACE2-binding affinities. At the same time, we noticed the reduced electrostatic contributions in BA.4/BA.5 which may be attributable to L452R and R493Q reversal mutations. A number of studies emphasized the role of electrostatic interactions as a dominant thermodynamic force leading at binding of the S-protein with the ACE2 receptor and antibodies.^114–116^ It was shown that the electrostatic potential surface of S protein and major variants to show accumulated positive charges at the ACE2-binding interface, revealing the critical role of complementary electrostatic interactions driving the enhanced affinity the Omicron S-ACE2 complexes.^115^

We compared the effect of the AF2-generated and MD-derived conformational ensembles on the MM-GBSA computations of binding affinities. Interestingly, both conformational ensembles yielded a strong correlation between the experimental dissociation constants and the predicted binding free energies (Figure 11A,B). The correlation graph of binding energies based on the AF2 conformational ensemble showed a larger dispersion of data points, while MD-based binding energies yielded only small standard deviations indicating that data points are clustered tightly around the mean. We also performed the residue-based decomposition of the MM-GBSA energies computed with MD ensembles. The analysis revealed that the binding energies are determined by contributions of only several hotspot centers (Figure 11C,D). The major contributors of binding affinity in the RBD-ACE2 complexes include RBD residues L455, F456, F486, N487, Y489, Q493, R498, T500, Y501 and G502. Among these sites, the strongest binding centers correspond to Omicron hotspots R498 and Y501 that are also known to cooperate in ACE2 binding via strong epistatic interactions.^46,47^ (Figure 11C,D). A comparison of the binding contributions for BA1, BA.2, BA.2.75 and BA.3 complexes indicated that the key hotspots at positions F486, Y489, F490, R493 (Q493), R498 and Y501 provide the most favorable contributions for BA.2 and BA.2.75 variants (Figure 11C). The second group of binding hotspots includes L455, F456 positions.

**Figure 11.**
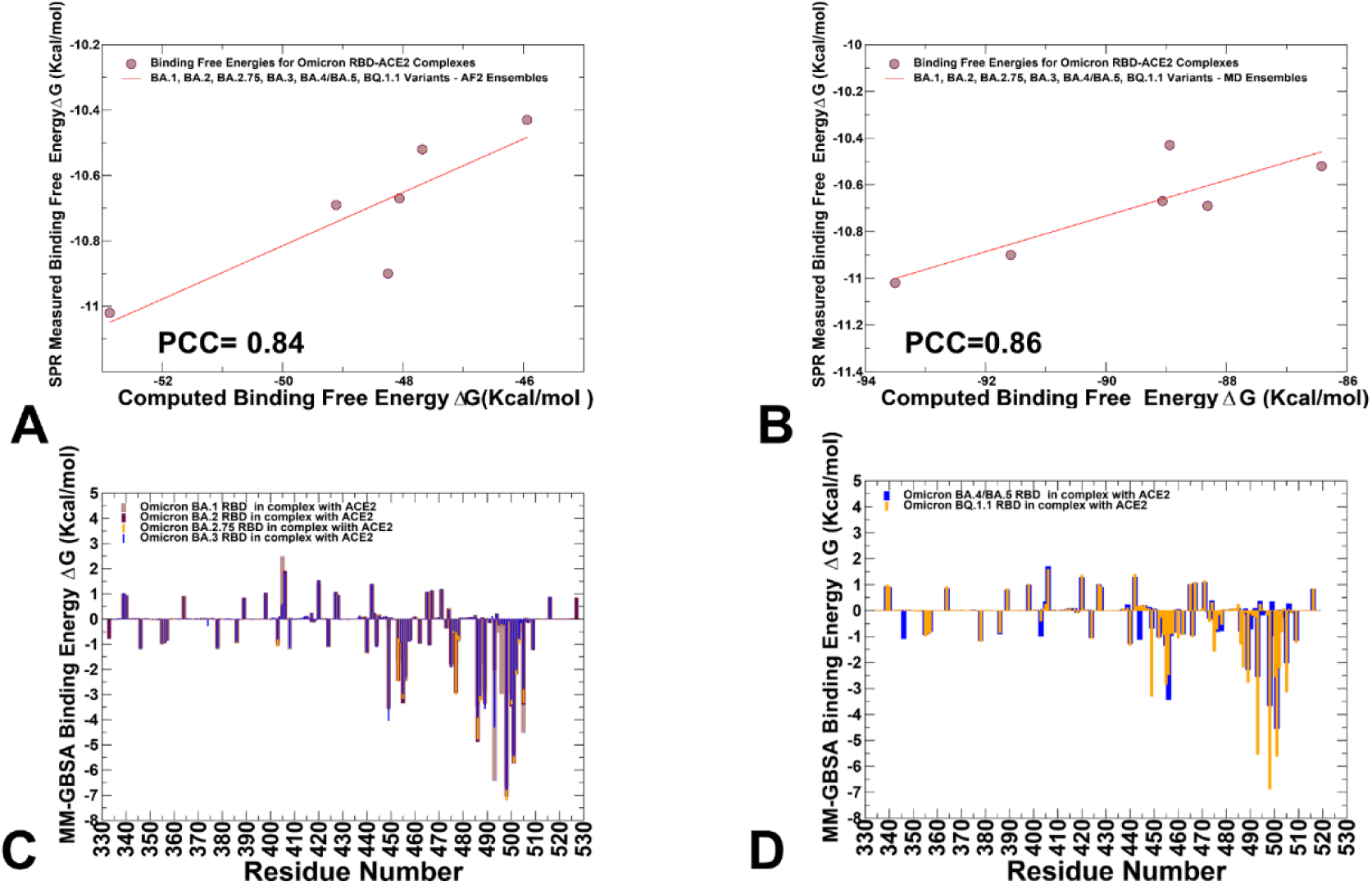
MM-GBSA binding energy analysis of the Omicron RBD-ACE2 complexes. (A) The scatter correlation graph between the computed binding free energies based on AF2-generated conformational ensembles and the SPR-measured binding affinities. (B) The scatter correlation graph between the computed binding free energies based on MD-generated equilibrium conformational ensembles and the SPR-measured binding affinities. The Pearson correlation coefficient values are indicated. (C) The residue-based decomposition of the total MM-GBSA binding energy ΔG contribution for the BA.1 RBD-ACE2 complex (light brown bars), BA.2 RBD-ACE2 complex (maroon bars), BA.2.75 RBD-ACE2 complex (orange bars) and BA.3 RBD-ACE2 complex (blue bars). (D) The residue-based decomposition of the total MM-GBSA binding energy ΔG contribution for the BA.4/BA.5 RBD-ACE2 complex (blue bars), and BQ.1.1 RBD-ACE2 complex (orange bars). The residue-based decomposition of the total MM-GBSA binding energies are based on the computations using MD equilibrium ensemble. The MM-GBSA contributions are evaluated using 1,000 samples from the equilibrium MD simulations of respective RBD-ACE2 complexes. The MM-GBSA calculations based on AF2 ensemble used 100 conformational states obtained from AF2-based shallow MSA subsampling and randomized sequence scanning combined with shallow MSA. It is assumed that the entropy contributions for binding are similar and are not considered in the analysis. The statistical errors for the residue-based decomposition of the total MM-GBSA binding energy ΔG contribution were estimated on the basis of the deviation between block average and are within 0.55-1.25 kcal/mol.

Mutational sites that contribute to the ACE2-binding affinity are also located in the flexible RBD regions, including D405N, N440K, L452R, S477N, T478K, E484A, Q493R, Q498R and N501Y, among which Q493R and N501Y are the most significant. In addition, small binding contributions provided by RBD residues K378, R403, K424, K440, K444, K460, N477, K478 are determined by strong electrostatic interactions mediated by lysine residues, which is the result of Omicron evolution leading to significant accumulation of positively charged substitutions interacting with the negatively charged ACE2 binding interface. To separately analyze differences between BA.4/BA.5 and BQ.1.1 complexes, we compared the contributions of individual RBD residues (Figure 11D). The breakdown highlighted stronger contributions of binding hotspots V486, Q493, R498 and Y501 in BQ.1.1 variant. Indeed, in BA.4/BA.5 the binding energy contribution for Q493 is ΔG = -2.54 kcal/mol while in the BQ.1.1 complex this contribution ΔG = -5.54 kcal/mol. Similarly, more favorable energy ΔG = -6.88 kcal/mol for Y501 in BQ.1.1 can be contrasted with less favorable ΔG = -4.54 kcal/mol for BA.4/BA.5 (Figure 11D). Of particular interest is comparison of contributions for convergent mutations in BQ.1.1 R346T, K444T, L452R, N460K, and F486V. It appeared that L452R and N460K sites are characterized by favorable electrostatic contributions resulting in the net ΔG =-1.5 kcal/mol for L452R and ΔG =-1.06 kcal/mol for N460K in BQ.1.1. In BA.4/BA.5 complex, we found a similar contribution of L452R position but the total binding contribution for N460 position is only ΔG =0.04 kcal/mol. At the same time, the presence of R346 and K444 in BA.4/BA.5 produces electrostatically-driven favorable contributions for both sites (ΔG = -1.08 kcal/mol for R346 and ΔG = -1.11 kcal/mol for K444). On the other side, R346T an K4454T mutations result in minor unfavorable contributions (ΔG = 0.18 kcal/mol for R346T and ΔG = 0.19 kcal/mol for K444T). The decreased charge on the BQ.1.1 RBD is due to mutations of two positively charged residues, R346T and K444T that only weakly interact with ACE2 which leads to a markedly reduced electrostatic contribution. These observations are consistent with previous studies showing that both R346T and K444T residues have a destabilizing effect on the interaction between RBD and ACE2.^117^ The negligible contributions of R346T and K444T mutational sites in BQ.1.1 were also confirmed in DMS study of BQ.1.1 variant in which mutations in these positions produced ΔG values within 0.1 kcal/mol and the reversed T346R and T444K modifications become basically neutral in the BQ.1.1 background.^52^ Hence, the emergence of reversed mutations introducing large positive chare appeared to produce a rather “muted” response in BQ.1.1 suggesting that electrostatic contribution becomes less dominant in this variant. In light of emergence of convergent mutations that reduce charge and may moderately reduce ACE2 binding, it may be speculated that the evolution of the virus is driven at least for BQ.1.1 variant by optimizing immune escape profile rather than its affinity for the receptor. Indeed, the latest study showed that the immune escape ability of BQ.1.1 can be attributed to R346T, K444T, and N460K modifications.^42^ Our results are consistent with this study showing that R346T and K444T are not involved in the interaction with ACE2, and these mutations do not perturb the loop structure compared to BA.4/BA.5. Another functional study demonstrated that R336T and K444T mutations may have emerged to reduce electrostatic interactions with antibodies and boost immune escape profile,^118^ suggesting that the observed reverse of the “electrostatic driving force” of ACE2 binding affinity for BQ.1.1 may be byproduct of optimized immune evasion profile.

Overall, the binding free energy computations established and confirmed several important findings. The central result of this analysis is the evidence that AF2-generated extended conformational ensembles can yield fairly accurate binding energies for Omicron RBD-ACE2 complexes leading to strong correlations with the experimental dissociation constants. Moreover, the observed correlations with the experiments using MD-derived equilibrium ensemble and AF2-generated conformational ensemble are quite similar. Hence, despite the lack of physically rigorous thermodynamic distribution of the equilibrium states in the AF2 predictions, AF2-based adaptations can efficiently explore and predict pools of functionally relevant conformational states that dominate the thermodynamics of the binding reaction. Moreover, our analysis offered support to the notion that using pLDDT assessment of structural confidence for screening of generated structures can efficiently determine the ensemble of functionally important spike conformations.

Another important finding of the binding energy analysis is that the effect of most Omicron RBD mutations on binding with ACE2 is rather small despite appreciable mutational modifications. Our results are consistent with the DMS experiments performed in different genetic backgrounds of BA,1, BA.2, BQ,1.1 and XBB.1.5 variants^52,120–122^ showing for example that many sites in the BA2 and BQ.1.1 RBDs are highly tolerant to mutations. These experimental studies showed that tolerance to mutations in many RBD positions is not a sign of insignificance as RBD residues that interact with ACE2 could also be involved in immune escape and exhibit considerable mutational plasticity. We observed the energetic dominance of binding energy hotspots R493 (or R493Q), R498, and Y501 across all studied Omicron variants. Moreover, a second group of important binding centers includes L455 and F456 central positions (Figure 11C,D). Importantly, the latest experimental evidence showed that ACE2 binding can be synergistically amplified via epistatic interactions of physically proximal binding hotspots, including Y501, R498, Q493, L455 and F456 residues.^52,120–122^ Hence, our results supported the notion that these epistasis-coupled RBD centers play a key role in convergent evolution by protecting ACE2 binding to compensate binding deficiencies of antibody-escaping mutations.^52^

The evolution of Omicron variants through acquisition of convergent mutational sites may leverage conformational adaptability and dynamic epistatic couplings between key binding hotspots to maintain and protect high ACE2 affinity while enabling the emergence of ACE2-neutral or partially destabilizing mutations that can modulate immune evasion.

## Conclusions

Atomistic level structural predictions and microsecond atomistic MD simulations provided a detailed characterization of the conformational ensembles and identified important differences in conformational landscape of the Omicron variants. Several different adaptations of the AF2 methodology along with MD simulations were used for a comparative characterization of structures, conformational ensembles and subsequent computations of binding affinities of the Omicron RBD-ACE2 complexes including BA.1, BA.2, BA.2.75, BA.3, BA.4/BA.5 and BQ.1.1 variants. By integrating predictions of the conformational states using AF2-based approaches we can efficiently expand characterization of the conformational ensembles for the RBD-ACE2 complexes capturing conformational details of the RBD fold and variant-specific functional adjustments of the RBD functional regions and mutational sites. We found that by evaluating the quality of the generated conformations using pLDDT metric and considering conformations with high confidence pLDDT values of ∼70-90 AF2 approaches can greatly expand coverage of the accessible conformational landscape and characterize functional protein ensembles. We leveraged AF2-based structural ensembles and MD-generated equilibrium ensembles for accurate comparative prediction of the binding energetics for the Omicron RBD-ACE2 complexes, which is consistent with the experimental data. The important finding of this analysis is that AF2-generated extended conformational ensembles can produce accurate binding energies for Omicron RBD-ACE2 complexes leading to strong correlations with the experimental dissociation constants. Our analysis offered support to the notion that using pLDDT assessment of structural confidence for screening of generated structures can efficiently determine the ensemble of functionally important conformations to compute binding affinities. The results suggested that combining MD simulations together with AF2-based prediction of conformational ensembles could provide more comprehensive view of the conformational landscapes and binding mechanisms for Omicron variants.

## Data Availability Statement

Data is fully contained within the article and Supplementary Materials. Crystal structures were obtained and downloaded from the Protein Data Bank (http://www.rcsb.org). All simulations were performed using the all-atom additive CHARMM36 protein force field that can be obtained from http://mackerell.umaryland.edu/charmm_ff.shtml. The rendering of protein structures was done with ChimeraX package (https://www.rbvi.ucsf.edu/chimerax/) and Pymol (https://pymol.org/2/). The software tools used in this study are freely available at the following GitHub sites : The software tools used in this study are freely available at GitHub sites : https://github.com/deepmind/alphafold; https://github.com/sokrypton/ColabFold/; https://github.com/RSvan/SPEACH_AF; https://www.github.com/HWaymentSteele/AFCluster; https://github.com/nextstrain; https://github.com/Amber-MD/cpptraj; https://github.com/smu-tao-group/protein-VAE; https://github.com/Amber-MD/cpptraj.

All the data obtained in this work including AF2-generated protein structures and conformational ensembles obtained using different approaches, simulation trajectories, topology and parameter files, the software tools, and scripts are freely available at ZENODO website https://zenodo.org/records/10904485.

## Author Contributions

Conceptualization, G.V.; methodology, N.R., M.A., G.C., S.X., G.V. P.T.; software, N.R., S.X., M.A., G.G., G.V and P.T.; validation, N.R., G.V.; formal analysis, N.R., G.V., M.A., G.G., S.X., and P.T.; investigation, N.R., M.A., G.C., G.V. and P.T.; resources, N.R., G.V., M.A. S.X., and G.V.; data curation, N.R., M.A., G.C.,G.V.; writing— original draft preparation, N.R., M.A., G.V.; writing—review and editing, N.R. and G.V.; visualization, N.R., M.A., G.C., S.X., G.V. G.V.; supervision, G.V.; project administration, G.V.; funding acquisition, P.T. and G.V. All authors have read and agreed to the published version of the manuscript.

## Conflicts of Interest

The authors declare no conflict of interest. The funders had no role in the design of the study; in the collection, analyses, or interpretation of data; in the writing of the manuscript; or in the decision to publish the results.

## Funding

This research was supported by the National Institutes of Health under Award 1R01AI181600-01 and Subaward 6069-SC24-11 to G.V. and National Institutes of Health under Award No. R15GM122013 to P.T.

## Acknowledgments

G.V acknowledges support from Schmid College of Science and Technology at Chapman University for providing computing resources at the Keck Center for Science and Engineering.

## Notes

### Competing Interest Statement

The authors have declared no competing interest.

