## Supplemental Figures S1-S5 for "Predicting Functional Conformational Ensembles and Binding Mechanisms of Convergent Evolution for SARS-CoV-2 Spike Omicron Variants Using AlphaFold2 Sequence Scanning Adaptations and Molecular Dynamics Simulations"

### Supplementary Materials

#### Exploring Conformational Landscapes and Binding Mechanisms of Convergent Evolution for the SARS-CoV-2 Spike Omicron Variant Complexes with the ACE2 Receptor Using AlphaFold2-Based Structural Ensembles and Molecular Dynamics Simulations


Nishank Raisinghani, [1] Mohammed Alshahrani,[1] Grace Gupta,[1] Sian Xiao[3], Peng Tao[3], Gennady Verkhivker[1,2]*

[1]Keck Center for Science and Engineering, Graduate Program in Computational and Data Sciences, Schmid College of Science and Technology, Chapman University, Orange, CA 92866, United States of America

[2] Department of Biomedical and Pharmaceutical Sciences, Chapman University School of Pharmacy, Irvine, CA 92618, United States of America

[3]Department of Chemistry, Center for Research Computing, Center for Drug Discovery, Design, and Delivery (CD4), Southern Methodist University, Dallas, Texas, 75275, United States of America

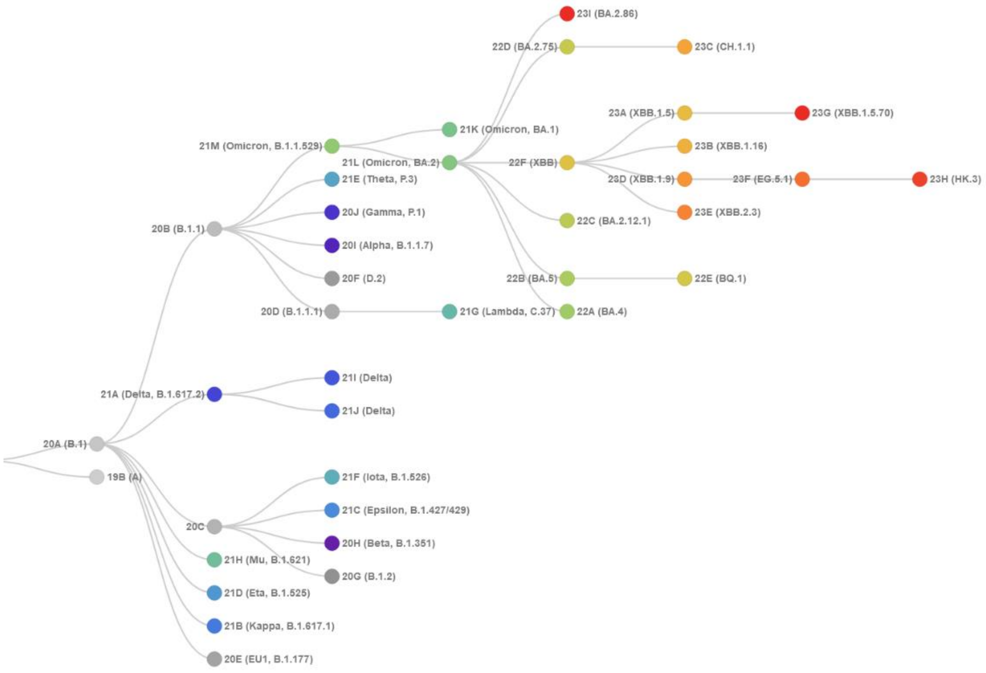


**Figure S1.** The evolutionary tree of current SARS-CoV-2 clades. XBB.1, XBB.1.5 and XBB.1.5.70 variants are shown on the current tree. The graph is generated using Nextstrain, an open-source project for real time tracking of evolving pathogen populations (https://nextstrain.org/).

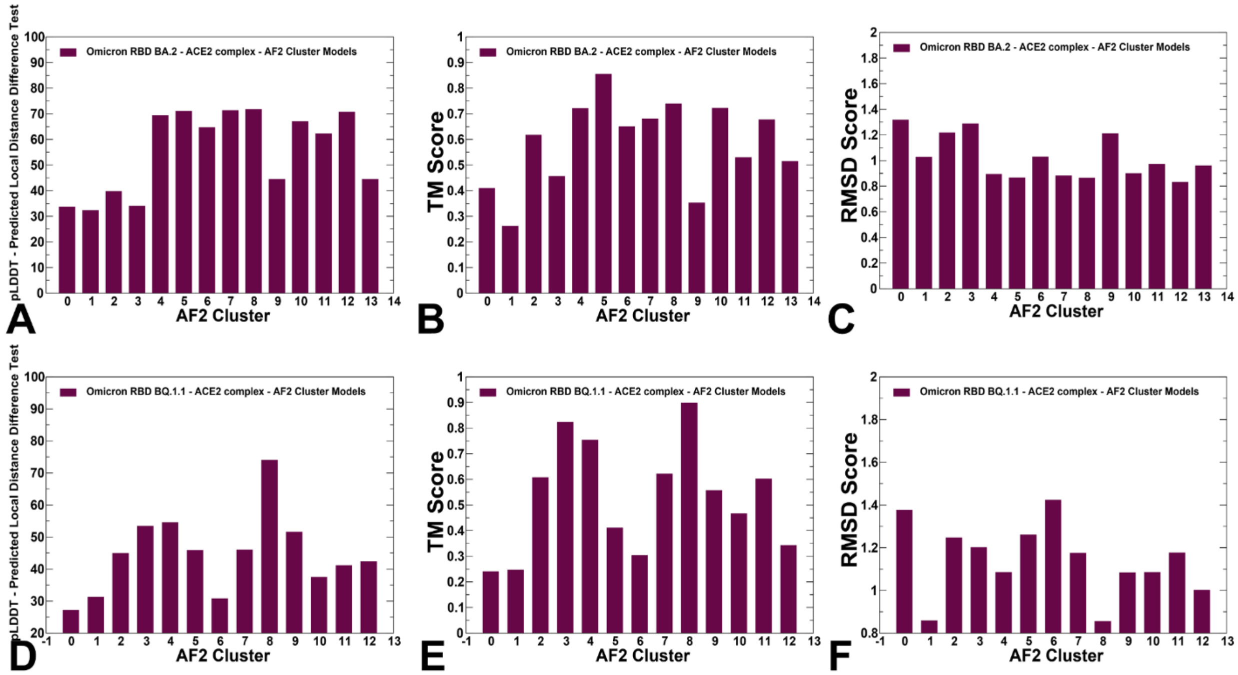


**Figure S2.** The distribution of statistical confidence pLDDT and structural similarity metrics for conformational cluster conformations obtained from AF-Cluster. The pLDDT profile as a function of cluster number from AF-Cluster predictions for BA.2 RBD-ACE2 complex (A) an BQ1.1 RBD-ACE2 complex (D). The TM-core profile as a function of cluster number from AF-Cluster predictions for BA.2 RBD-ACE2 complex (B) an BQ1.1 RBD-ACE2 complex (E). The RMSD from the experimental structure profile as a function of cluster number from AF-Cluster predictions for BA.2 RBD-ACE2 complex (C) an BQ1.1 RBD-ACE2 complex (F).

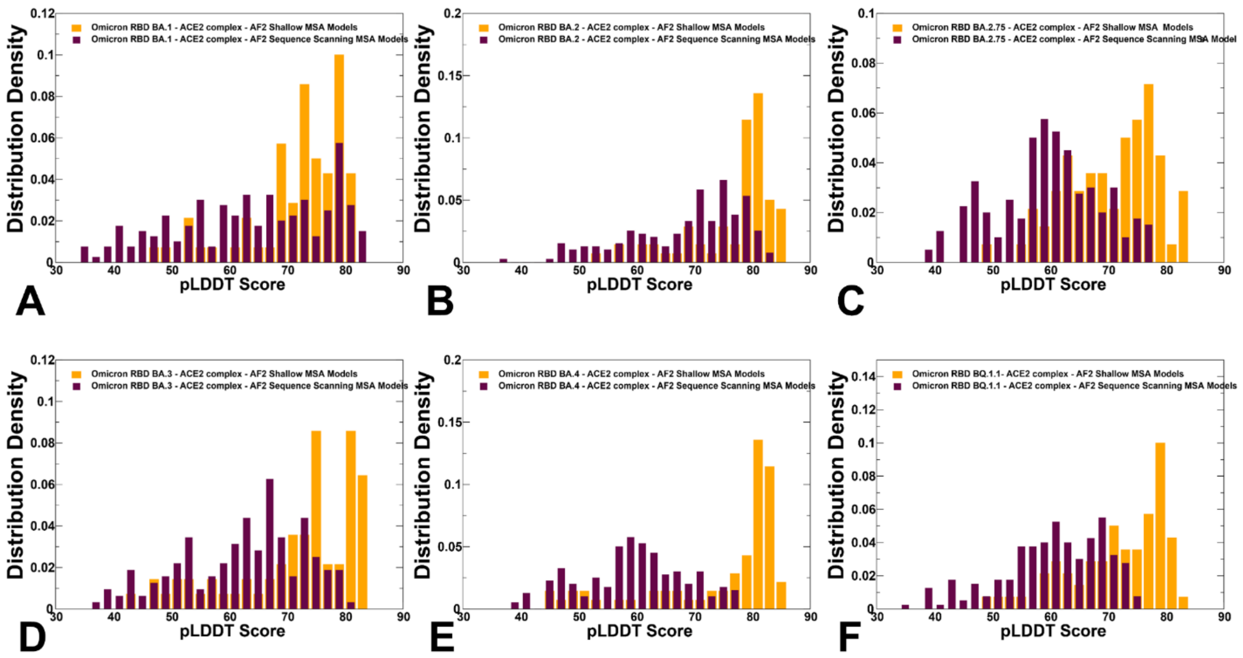


**Figure S3.** The distributions of the pLDDT metric for the RBD-ACE2 conformational ensembles obtained from AF2-MSA shallow depth predictions and AF2 with randomized sequence scanning. The density distribution of the pLDDT values for structural ensembles are shown for BA.1 (A) BA.2 (B), BA.2.75 (C), BA.3 (D), BA.4/BA.5 (E) and BQ.1.1 (F). The pLDDT density distribution for conformations obtained using AF2-MSA shallow depth are shown in orange-colored filled bars and the pLDDT distribution for conformations using AF2 with randomized sequence scanning are in maroon filled bars.

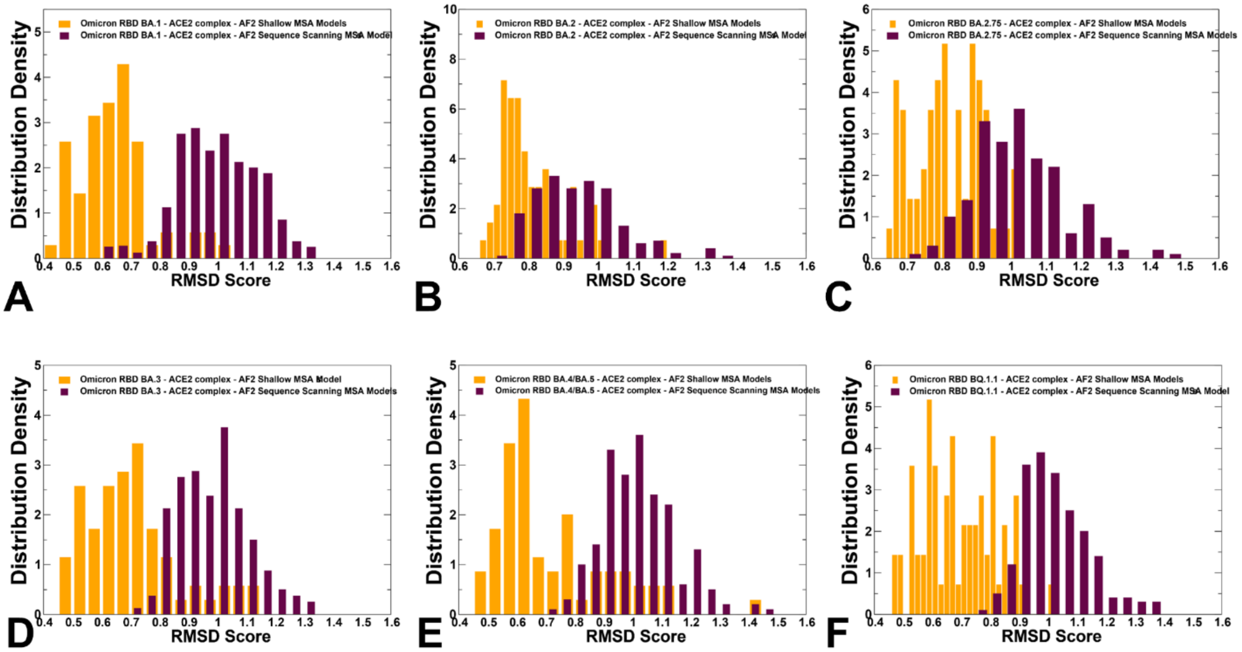


**Figure S4.** The distributions of the RMSD values from the experimental structures for the RBD-ACE2 conformational ensembles obtained from AF2-MSA shallow depth predictions and AF2 with randomized sequence scanning. The density distribution of the RMSD values for structural ensembles are shown for BA.1 (A) BA.2 (B), BA.2.75 (C), BA.3 (D), BA.4/BA.5 (E) and BQ.1.1 (F). The RMSD density distribution for conformations obtained using AF2-MSA shallow depth are shown in orange-colored filled bars and the RMSD distribution for conformations using AF2 with randomized sequence scanning are in maroon filled bars.

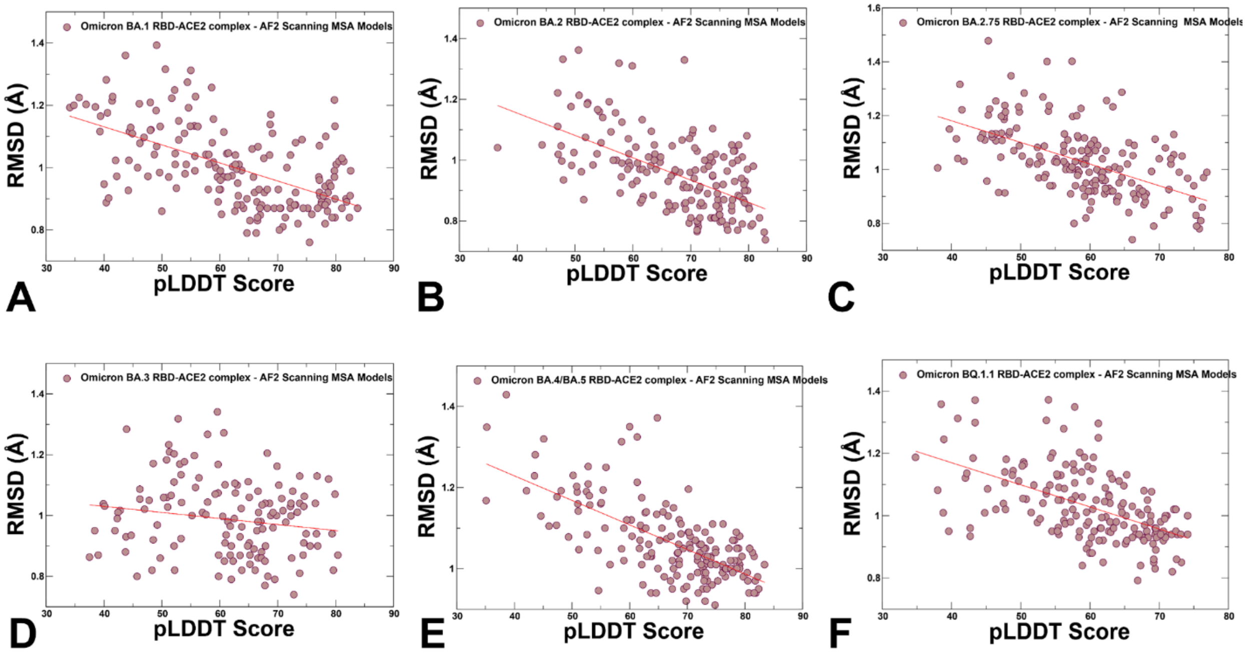


**Figure S5.** The scatter plots between the pLDDT scores and RMSD scores for the conformational ensembles obtained using AF2 adaptation with randomized sequence scanning and subsequent shallow MSA subsampling. The scatter plot distributions are shown for structural ensembles obtained for BA.1 (A) BA,2 (B), BA.2.75 (C), BA.3 (D), BA.4/BA.5 (E) and BQ.1.1 (F). The density distributions are depicted as brown-colored filled circles.
